# ChIPulate : A comprehensive ChIP-seq simulation pipeline

**DOI:** 10.1101/467241

**Authors:** Vishaka Datta, Sridhar Hannenhalli, Rahul Siddharthan

## Abstract

ChIP-seq (Chromatin Immunoprecipitation followed by sequencing) is a high-throughput technique to identify genomic regions that are bound in vivo by a particular protein, e.g., a transcription factor (TF). Biological factors, such as chromatin state, indirect and cooperative binding, as well as experimental factors, such as antibody quality, cross-linking, and PCR biases, are known to affect the outcome of ChIP-seq experiments. However, the relative impact of these factors on inferences made from ChIP-seq data is not entirely clear. Here, via a detailed ChIP-seq simulation pipeline, ChIPulate, we assess the impact of various biological and experimental sources of variation on several outcomes of a ChIP-seq experiment, viz., the recoverability of the TF binding motif, accuracy of TF-DNA binding detection, the sensitivity of inferred TF-DNA binding strength, and number of replicates needed to confidently infer binding strength. We find that the TF motif can be recovered despite poor and non-uniform extraction and PCR amplification efficiencies. The recovery of the motif is however affected to a larger extent by the fraction of sites that are either cooperatively or indirectly bound. Importantly, our simulations reveal that the number of ChIP-seq replicates needed to accurately measure in vivo occupancy at high-affinity sites is larger than the recommended community standards. Our results establish statistical limits on the accuracy of inferences of protein-DNA binding from ChIP-seq and suggest that increasing the mean extraction efficiency, rather than amplification efficiency, would better improve sensitivity. The source code and instructions for running ChIPulate can be found at https://github.com/vishakad/chipulate.

## Introduction

ChIP-seq (Chromatin immunoprecipitation and sequencing) is a popular high-throughput experimental technique to find locations that are bound *in vivo* by a single transcription factor (TF) [1]. Upon mapping of the DNA fragments bound by the TF to the reference genome, the genomic loci bound by the TF are identified as high density mapped regions or *peaks*, where each peak is associated with an *intensity* based on the number of sequenced fragments arising from it. The intensity reflects the *in vivo* occupancy of the TF at that locus.

Several studies of ChIP-seq data have focussed on the biological factors distinguishing the loci bound by the TF. It has been shown that in addition to the affinities of binding sites present at a locus, nucleosome positioning is a strong determinant of TF binding *in vivo* [2, 3, 4, 5]. Other studies have shown that the concentration of the target TF [6, 7], short-range cooperative interactions between the target TF and other TFs [8], and variation in chromatin accessibility [5, 7] explain the variation in intensities across peaks. Some of the variation can arise due to indirect binding, where the target TF binds DNA indirectly via a second DNA-bound TF [9, 10, 11]. The intensity of such peaks is then no longer directly dependent on the affinity of the target TF to sequence at the bound locus.

Since the distribution of ChIP-seq peaks and their intensities depend on factors other than the affinity of the target TF towards sequence at a locus, it impacts two kinds of inferences often made from ChIP-seq data. First, the highest intensity peaks are used to infer position weight matrix (PWM) or motif models of the target TF [12]. While it is known that changes in the concentration of the target TF can distort the inferred PWM [13, 14], the extent to which cooperative and indirect interactions distort the PWM is unclear. Second, along with variations in chromatin accessibility, these interactions weaken the statistical dependence of peak intensity on target TF binding site affinity, which means that a peak with a higher intensity need not necessarily contain a higher affinity target TF binding site.

In addition to these biological factors, ChIP-seq peaks are affected by purely experimental sources of noise. The ChIP-seq protocol broadly consists of three key steps [1, 15, 16] — (i) the extraction of fragments that are bound by the target TF (ii) PCR amplification of these extracted fragments, and (iii) sequencing of these fragments. It is known that fragments can be extracted more easily at some genomic regions than others due to differences in chromatin structure and cross-linking efficiency across the genome [17, 18, 19]. Similarly, certain fragments are more efficiently amplified by PCR than others due to differences in GC content or the presence of nucleotide repeats [20, 21, 22]. The extent to which these factors distort peak intensities and make otherwise identical genomic regions appear differentially occupied has not been quantified. Multiple biological replicates of ChIP-seq are recommended [23] to overcome these issues, but the quantitative improvement in accuracy as more replicates are performed has not been ascertained.

To evaluate the influence of the aforementioned biological and experimental sources of variation on ChIP-seq peak intensities, we have developed a comprehensive pipeline to simulate a ChIP-seq experiment called ChIPulate. In particular, we simulate genome-wide TF-DNA binding based on established biophysical models of binding and also simulate the extraction, amplification and sequencing steps of the ChIP-seq protocol. Our simulation thus associates a set of read counts at each genomic locus with its occupancy by the target TF. Whereas existing statistical models [24, 25, 26, 14] assume that read counts follow a prescribed probability distribution and treat the ChIP-seq protocol as a black box, we explicitly model key steps of the ChIP-seq protocol. Our approach allows us to individually evaluate the impact of variation in extraction and PCR amplification efficiencies, as well as chromatin accessibility, indirect binding, and cooperative binding, on peak intensities and PWM inference. We are also able to quantify the extent to which additional replicates of ChIP-seq improve its ability to robustly measure the occupancy of a genomic region.

We find that biological factors such as indirectly and cooperatively bound sequences distort inferred PWMs more than experimental sources of variation. Variations in extraction efficiency across the genome distort peak intensities and lower their ability to distinguish between occupied loci. Poor extraction efficiency also increases the number of false positive peaks, which are peak calls that do not contain a binding site for the target TF. In contrast to the effect of variations in extraction efficiency, even drastic variations in PCR amplification efficiency have little impact on peak intensities, and hence do not affect the inferred PWM or increase the number of false positive peaks. Finally, we found that at least two biological replicates of ChIP-seq read counts are necessary to reliably infer the binding energy of a genomic region.

Our work provides a general framework and a software tool for simulating ChIP-seq read counts through a realistic model of TF-DNA binding and the steps of the ChIP-seq protocol. Improvements in the protocol to lower variation in extraction efficiency, rather than PCR amplification efficiency, are more likely to improve our ability to distinguish between genomic regions of differing occupancy. Further, changes in the protocol that allow multiple biological ChIP-seq replicates to be performed, or computational approaches that can reliably combine read counts from ChIP-seq experiments, would allow more accurate inferences of TF-DNA occupancy, especially in regions containing multiple binding sites.

## Results

### A framework to simulate read counts from TF binding sites in a ChIP-seq experiment

A schematic of our ChIP-seq simulation framework is shown in Figure 1A, with details in Methods. Broadly, the goal of our simulation is to take as an input the occupancy of a TF at multiple locations across a genome and output a set of read counts in both ChIP and input experiments at each location. A single TF’s occupancy at a genomic location is determined by its site-specific binding energy, and its chemical potential, which depends on the logarithm of the concentration of the TF [27].

**Figure 1:**
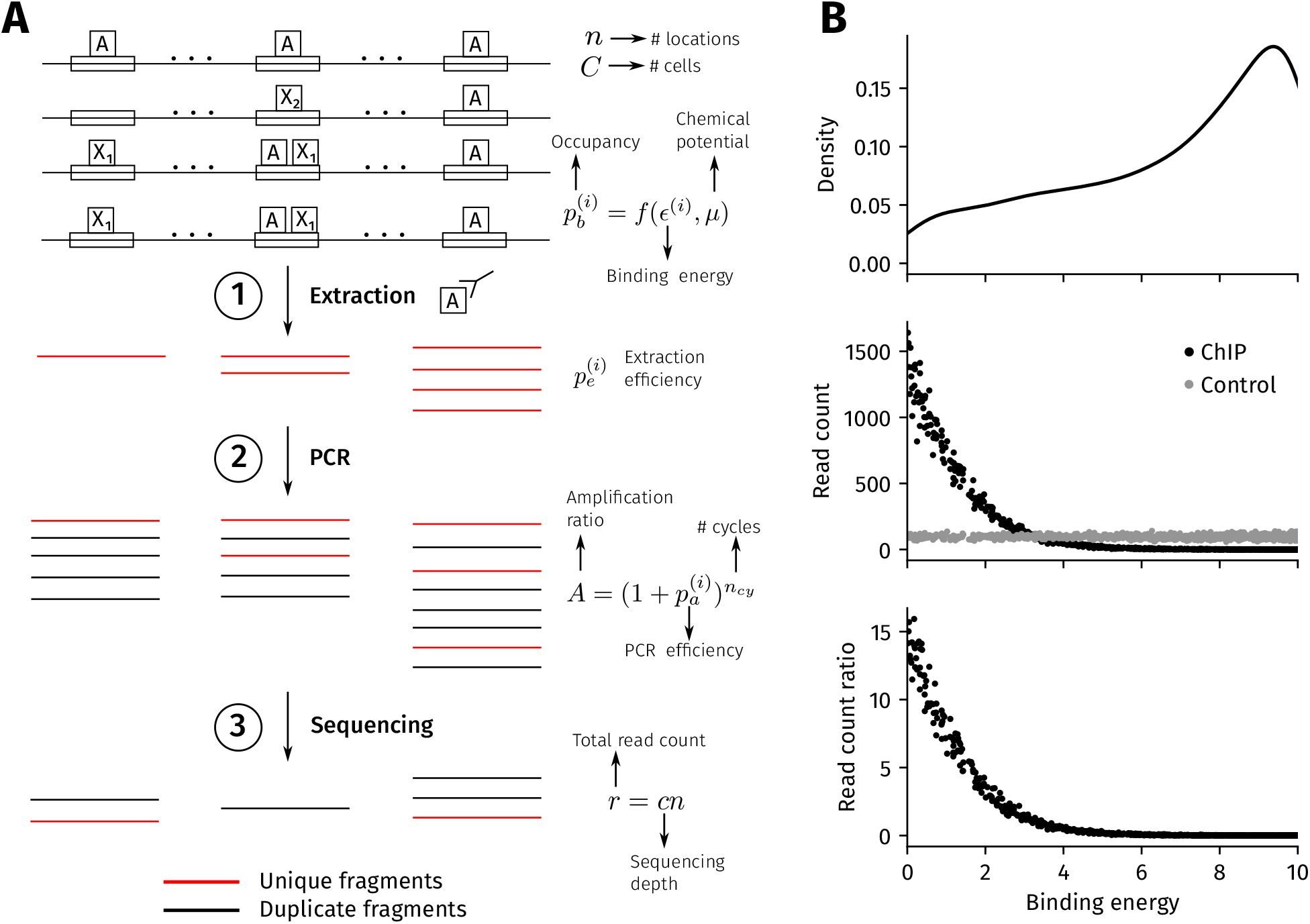
ChIP-seq simulation procedure. **(A) Procedure overview**. Each of *n* genomic regions contains a binding site with a binding energy *ϵ*^(*i*)^ (in units of *k_B_T*), where a lower value represents a higher affinity binding site. In the ChIP sample, the binding energy at the i-th location, and the chemical potential, set the probability of occupancy 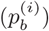 of the target TF. The probability of occupancy determines the number of cells (out of a total of *C* cells) where the i-th location is bound. In Step 1 of the simulation, a total of 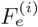 fragments are extracted at the i-th location from a binomial distribution with mean 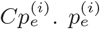 is the extraction efficiency that represents the probability of successfully extracting a bound fragment. All of these fragments are unique (in red) since each fragment originates from a different cell. In Step 2, the extracted fragments are amplified through *n_cy_* cycles of PCR to give 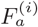 amplified fragments (duplicate fragments in black) with the probability of amplification of each fragment, or PCR efficiency, being 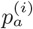 in every cycle. The average number of amplified fragments A obtained from each extracted fragment is 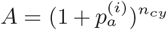, which we refer to as the amplification ratio. In Step 3, *r_i_* reads are obtained from the i-th location after sampling *r = cn* fragments from all 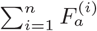 amplified fragments, where *r* is the total read count and *c* is the sequencing depth. The number of unique reads at each location is the final output of the simulation. For the control input sample, read counts are simulated as for the foreground ChIP sample, except that the probability of occupancy is assumed to be constant at all *n* locations and the number of cells is assumed to be 0.1*C*. **(B) Sample output from the simulation**. The top panel shows the distribution of binding energies employed in the simulation, which by default is a truncated power law between 0 and 10*k_B_T* with an exponent of 0.5. The middle panel shows the number of unique reads from each genomic region in the ChIP (black) and input (gray) experiments. The bottom panel is the read count ratio between the ChIP and input samples at each location. The simulation parameters are *n* = 1000, *μ* = 3*k_B_T*, *ϵ_bg_* = 1*k_B_T*, *C* = 10^5^, *n_cy_* = 15, and *A* = 1000. The extraction efficiencies in the ChIP and input samples follow a truncated normal distribution between 0 and 1, with a mean of 0.5 and standard deviation of 0. 05 across the genome.

Each genomic location is associated with two experimental parameters — its extraction efficiency and PCR efficiency. The extraction efficiency, which determines how many bound fragments are successfully extracted in Step 1 in Figure 1A, can vary between different genomic locations. The extracted fragments are then subjected to a number of PCR amplification cycles in Step 2. The average number of amplified fragments obtained at each location depends on the PCR efficiency associated with the locus, which can also vary between different locations. Step 3 represents the sequencing step where the amplified fragments from all locations are sequenced. The number of sequenced fragments is the total read count of the experiment. Since unique and duplicate fragments (red and black in Fig. 1A) are tracked through the simulation, the exact number of unique reads at each location is known.

In addition to the ChIP experiment, we simulate the sequencing of a genomic control input sample. This simulation differs from the ChIP experiment in two ways. First, the background binding energy at each occupied genomic locus is set to a fixed value of 3*k_B_T* for all locations (see Methods for justification). Second, we set the number of cells to be only 10% of the number of cells employed in the ChIP simulation. A larger number of cells are used in the ChIP simulation since, in practice, DNA is less efficiently extracted in the ChIP sample due to the use of an antibody whereas no antibody is used in the input sample. Thus, to extract a given amount of DNA, fewer cells are employed in the input sample than in the ChIP sample. In the event that the number of extracted fragments in the input is more than the number of extracted fragments in the ChIP sample, we down-sample the input fragments such that the total number of extracted fragments is identical in both ChIP and input samples before the PCR amplification step. Similarly, we down-sample the ChIP fragments if the number of extracted fragments in the ChIP sample is more than than that in the input sample. At each locus, the extraction and PCR efficiencies in the input are kept identical to the values used in the ChIP simulation.

After simulating both ChIP and input experiments, we compute the ratio of the read counts obtained from the ChIP sample to that of the input sample at each genomic locus. The read count ratio at a location is equivalent to peak intensity computed by commonly used peak callers from ChIP-seq reads [24]. The read counts obtained from our simulations are shown in Figure 1B with the values of the simulation parameters set to the default values described in Methods.

### Key assumptions

When simulating genome-wide TF-DNA binding before Step 1 in Figure 1A, we assume that there is a single TF and a single binding site in each genomic region. We relax this assumption in later sections when simulating indirect and cooperative binding. We assume that binding in different genomic locations are independent of each other.

The extraction efficiency parameter in Step 1 aggregates the effect of the many steps of fragmentation, cross-linking, pull down and size selection that occur in a ChIP-seq experimental protocol. We show in Methods that this parameter takes all these steps into account when modeling the extraction of DNA fragments from cells. In the PCR amplification step, we assume the amplification process to be in the exponential phase of amplification [28] where increasing the number of PCR cycles exponentially increases the number of amplified fragments. We assume that any potential PCR mutational errors do not change the amplification ratio at any genomic location. In the sequencing step 3, we assume that the sequencing and mapping are error-free, i.e., reads are uniquely and correctly mapped, and they can be perfectly de-duplicated at each location.

### Motif inference is not affected by extraction and PCR efficiency whereas fidelity at locations with low read count ratios is affected by a low extraction efficiency

We evaluated the impact of heterogeneity in extraction and PCR efficiencies on two outcomes of ChIP-seq. One is the recoverability of TF motif based on the top genomic locations in terms of their read count ratios. The second is fidelity, defined as the probability (frequency) with which a site X with a read count ratio that is at least 10% higher than site Y in fact contains a sequence with a higher affinity (or lower binding energy).

We simulated the process of inferring the motif of an arbitrarily chosen *S. cerevisiae* transcription factor (Tye7) from ChIP-seq (Figure 2A). We assigned binding energies for 1000 genomic regions from our default binding energy distribution, which is a power law distribution (with an exponent of 0.5) between 0 and 10*k_B_T* (see Methods). By using the Tye7 binding energy matrix estimated by BEEML from protein binding microarray measurements [29, 13], we found the 10 bp sequence whose binding energy was closest to this assigned value, and virtually planted it at each location. We simulated a ChIP-seq experiment with a fixed extraction and PCR efficiency across the genome, following which we selected sites having the top 100 read count ratios and constructed the PWM of Tye7. This PWM, which we refer to as the *baseline PWM*, is used to compute the extent to which heterogeneity in extraction and PCR efficiency changes the derived PWM.

**Figure 2:**
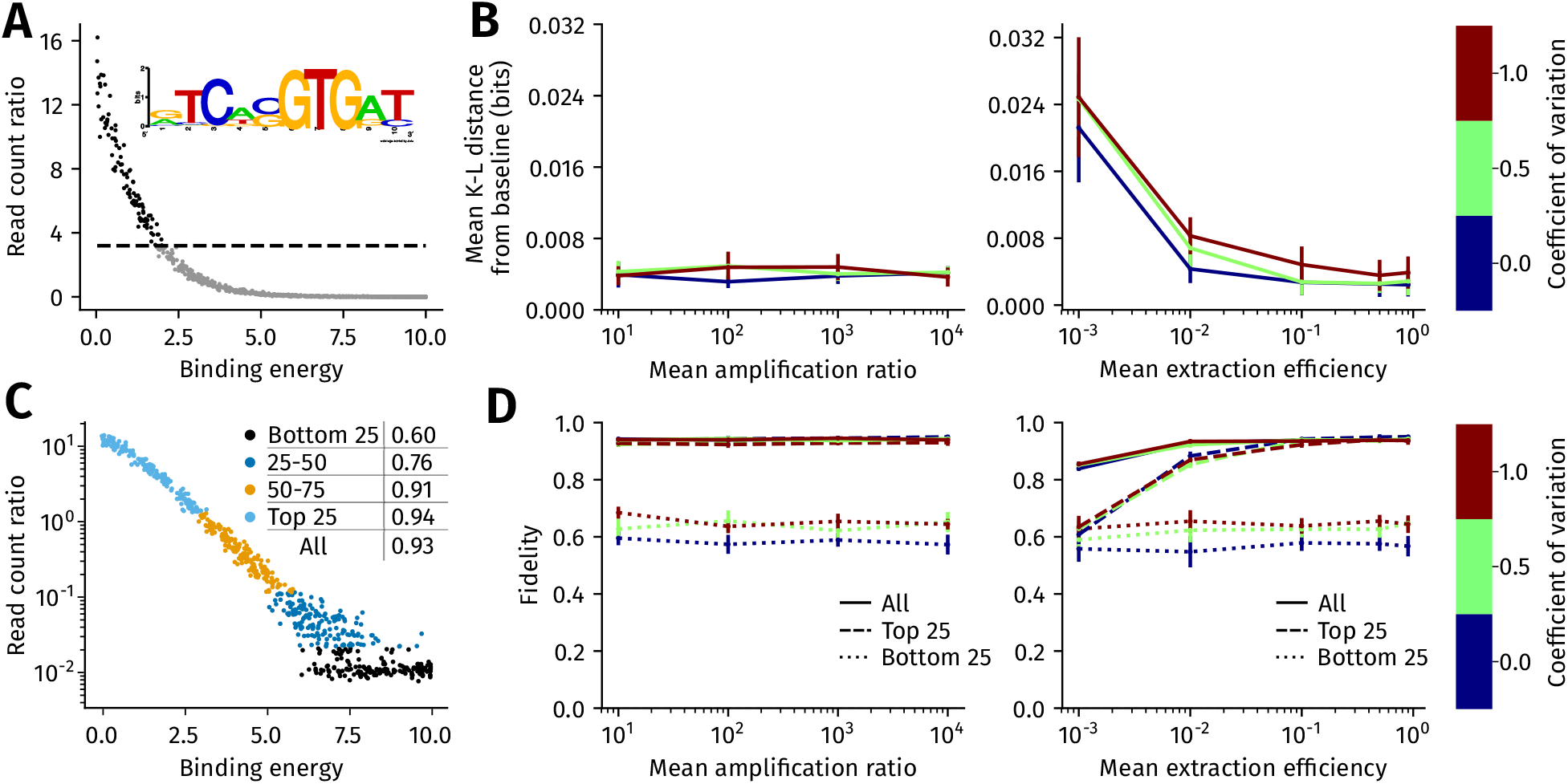
Impact of genome-wide heterogeneity in extraction and PCR efficiency on motif inference and ChIP-seq fidelity. **(A)** To simulate motif inference, 1000 binding energies were sampled from the default binding energy distribution. A binding site sequence of transcription factor Tye7 was assigned to each binding energy (Methods). After simulating ChIP-seq in the absence of extraction and amplification ratio heterogeneity, binding sites from locations with the 100 highest read count ratios were used to construct a baseline PWM of Tye7 (shown inset). **(B) The mean K-L distance between the baseline PWM and the motif inferred in the presence of heterogeneity in extraction efficiency (right) and amplification ratio (left)**. The heterogeneity in extraction and amplification ratio is assumed to follow a truncated normal distribution, with the mean increasing from left to right on the x-axis in both panels. The coefficient of variation of the truncated normal varies from 0 (no variation, in blue) to 0.5 (green) and 1.0 (brown). The error bars are the standard deviation in the mean K-L distance computed after PWM was estimated in 10 replicates of ChIP-seq for each mean and coefficient of variation. **(C) ChIP-seq fidelity captures the monotonicity of the relationship between read count ratio and binding energy**. Fidelity is defined as the probability that if a location i has a read count ratio at least 10% higher than location *j*, then it implies that *i* has a lower binding energy than *j*. Fidelity is calculated by sampling 1000 pairs of locations, where each pair could be from anywhere in the genome or top 25th, 25-50th,50-75th, or bottom 25th percentiles of the read count ratio. Read count ratios falling in each percentile bin are marked in different colors in the scatter plot, where the y-axis is plotted on a logarithmic scale. The fidelity values in each of these bins is shown in the plot legend, along with the fidelity computed across all regions. **(D) Variation in ChIP-seq fidelity with heterogeneity in extraction efficiency (right) and PCR amplification ratio (left)**. The x-axis and the three plot colors are defined identically to (B). The error bars are the standard deviation in the estimate of fidelity, which are computed from 10 replicates of simulation for a given mean and coefficient of variation.

Ideally, one would use a PWM of Tye7 published in a database as the baseline motif against which to make these comparisons. However, these motifs can differ between databases due to differences not only in the experimental assay used to measure TF-DNA binding but also the underlying binding energy distribution of the bound sequences. This means that the baseline Tye7 PWM derived from the simulation may differ from that present in the ScerTF database [30]. However, our goal is to specifically assess the effect of extraction and PCR amplification heterogeneity on motif recoverability. For this purpose, it suffices to compare the motif derived in the presence of heterogeneity with the baseline motif derived in the absence of heterogeneity.

Figure 2B shows the difference between the baseline PWM and the Tye7 PWM that is derived in the presence of extraction or PCR amplification heterogeneity, respectively, when the efficiencies follow a truncated normal distribution. This difference is measured in terms of the mean Kullback-Leibler (K-L) distance, measured in bits (Methods), between the two PWMs. The impact of heterogeneity in these parameters on the inferred motif is small as the highest K-L distance from the baseline motif is about 0.045 bits. The effect of PCR amplification and extraction heterogeneity on motif inference is more pronounced when the heterogeneity follows a power law distribution (Supplementary Figure 1A) than when it follows a truncated normal distribution. The distortion of the PWM is still relatively low, however, with the highest K-L distance from the baseline motif being less than 0.1 bits per position. We also evaluated the impact of varying the chemical potential between 1*k_B_T* and 6*k_B_T* on the inferred motif when the baseline motif is derived at *μ* = 3*k_B_T*. Examples of the effect of a different chemical potential on the read count ratios obtained across the genome are shown in Supplementary Figure 2A. We found that the inferred motif deviates up to 0.08 bits per position from the baseline motif (Supplementary Figure 2B). This is within the same order of magnitude as the effect of a low mean extraction efficiency on the inferred motif.

We checked the extent to which the trends in Figure 2B depended on our choice of TF. We selected 21 TFs that possessed different DNA binding domains [31] (Supplementary Table 1), and for each TF, we simulated motif inference when there is heterogeneity in extraction efficiency (Supplementary Figure 3) or the PCR amplification ratio (Supplementary Figure 4) across the genome. The binding energy distribution of each TF, along with all other parameters of ChIPulate, was kept at their default values. We found that motif inference in each of these TFs was affected in a manner similar to Tye7, where a low mean extraction efficiency distorts the inferred motif more than a low mean amplification efficiency. However, we found that the extent of motif distortion due to extraction heterogeneity was significantly correlated (*R*^2^ = −0.67, *p* < 10^-3^) with the motif information content of each TF (Supplementary Figure 3A), while there was no such correlation in the presence of amplification heterogeneity (Supplementary Figure 4A, *R*^2^ = −0.36, *p* = 0.1). Finally, we found a weak dependence of the extent of motif distortion on the length of the motif, where a low mean extraction efficiency distorted a 6 bp motif upto nearly 0.1 bits per position with a lower magnitude of distortion for longer motifs (Supplementary Figure 5A). There was no such dependence of motif distortion on motif length when the amplification ratio varied across the genome (Supplementary Figure 5B).

Results shown in Figure 2C show that ChIP-seq fidelity is generally higher for sites with intermediate read count ratios (50th percentile to 75th percentile) and deteriorates at lower and higher read count ratios. This is partly due to the value of the chemical potential of 3*k_B_T* employed in our simulation. When the chemical potential is varied between 1*k_B_T* and 6*k_B_T*, the fidelity amongst the top 25th percentile of read count ratios noticeably decreases and reaches close to 0.5 at μ= 6*k_B_T* (Supplementary Figure 2C). The fidelity amongst read count ratios of the bottom 25%, however, increases over the same range of the chemical potential. This is due to the fact that the occupancy of locations whose binding energies are much lower than the chemical potential are close to 1 while locations with binding energies above the chemical potential are close to zero. As the chemical potential increases, the increase in occupancy at locations with high binding energies is sufficiently large for their read count ratios to differ from one another. On the other hand, the occupancies of locations with low binding energies rapidly tend to one, which results in similar read count ratios between sites that possess different binding energies.

Figure 2D shows the impact of heterogeneity in extraction efficiency and PCR amplification ratio on fidelity. The fidelity is highest when computed across all binding site pairs, than amongst pairs in the top 25% of read count ratios. This is because the expected difference between the binding energies of sites in each pair is larger when the pairs are selected from the entire range. Changes in the mean and variance in the amplification ratio have little impact on fidelity, both overall as well as in each read count ratio bin. The fidelity across all regions is also not substantially affected by the mean and variance in extraction efficiency, but the fidelity is markedly lowered for sites with the lowest read count ratios. When PCR amplification heterogeneity is power-law distributed, its impact on fidelity is still quite low (Supplementary Figure 1B). In contrast, when the extraction efficiency is power-law distributed across the genome, its impact on fidelity is more drastic than when it is normally distributed.

### A low mean extraction efficiency increases the probability of false positive peak calls but the PCR efficiency has no impact

We tested the effect of heterogeneity in extraction and PCR efficiency on the ability of ChIP-seq to distinguish peak calls that contain a binding site for the target TF (true positive) from those peak calls that do not contain a binding site for the target TF (false positive). Figure 3A shows a simulation where genomic locations that do not harbor a binding site for the target TF give rise to sequence reads. We refer to such locations as false positive peaks. This differs from indirectly bound peaks, where the target TF is part of the complex of TFs that binds DNA. In practice, a false positive peak can arise due to poor antibody quality, where the antibody binds epitopes on TFs other than the target TF. We assign binding energies to these false positives locations from a truncated power law such that the mean occupancy ratio between true and false positive sites are fixed while the variance in binding energies amongst false positive sites and true positive sites are equal. Figure 3B shows the receiver operating characteristic (ROC) curve for the simulation shown in Figure 3A. The ROC curve shows the change in the number of true positives and false positives when the read count ratio threshold at which a location is declared as a peak is changed. The area under the ROC curve (auROC) provides an overall measure of the accuracy of using ChIP-seq read count ratios in distinguishing between locations with target TF binding sites from those without these sites.

**Figure 3:**
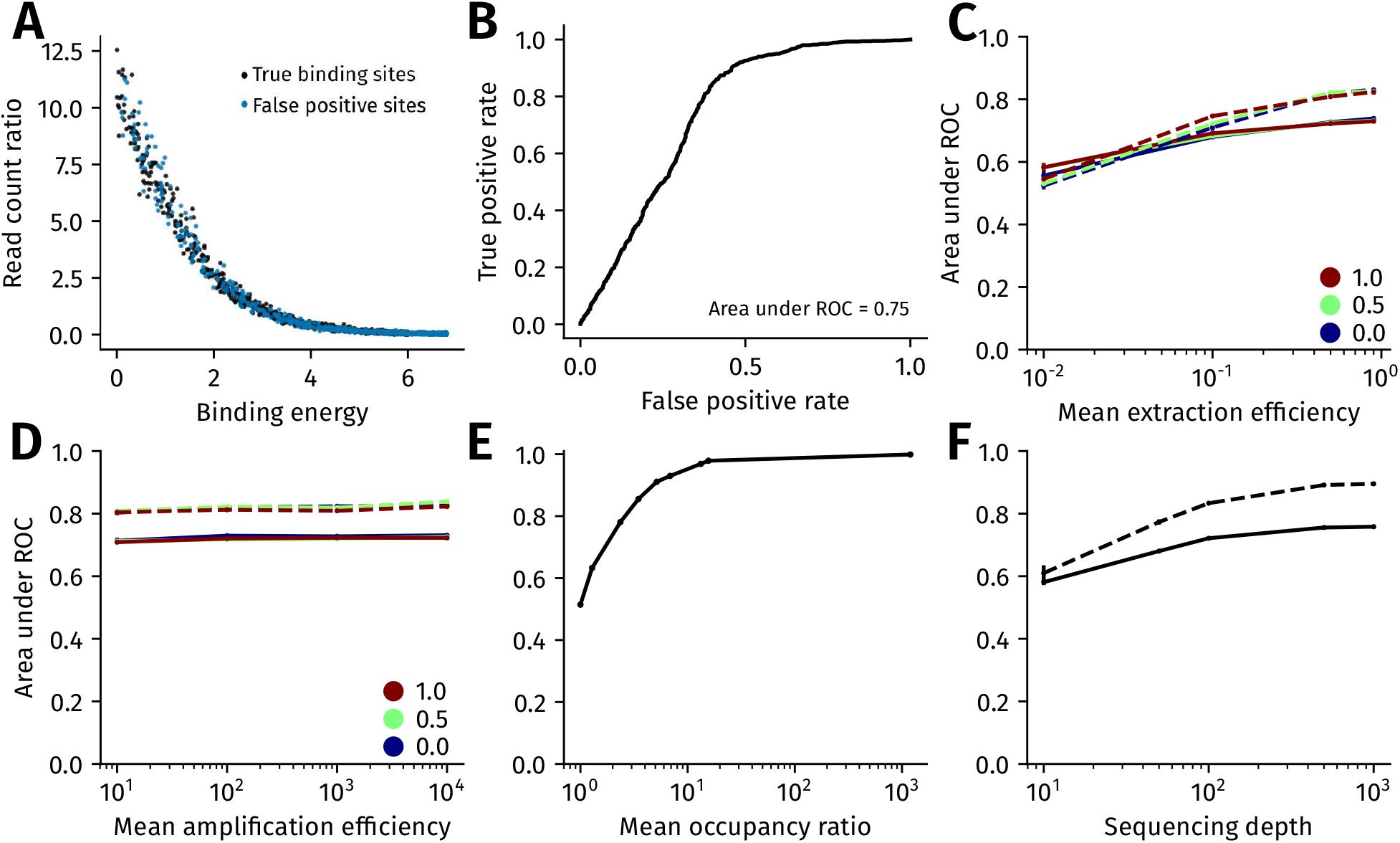
The impact of experimental and biological sources of variation on the sensitivity of ChIP-seq. **(A)** Simulating false positive binding sites. The binding energies of false positive genomic locations, which do not contain a target TF binding site, are distributed according to a truncated power law with exponent 0.76 in the range [0, 6.78*k_B_T*]. Binding energies of true positive genomic locations, which contain a target TF binding site, are sampled from a truncated power law with exponent 0.5 in the range [0, 6*k_B_T*]. **(B) Receiver operating characteristic (ROC) curve corresponding to the simulation shown in A. (C,D) Variation in auROC with the extraction efficiency and PCR amplification ratio**. The mean extraction efficiency in (C) and the mean amplification efficiency (after 15 cycles of PCR) in (D) increases along the x-axis. The efficiencies vary according to a truncated normal distribution with the blue, green and brown lines corresponding to a coefficient of variation of 0, 0.5 and 1.0, respectively. The solid and dashed lines are the auROC when the ratio of the mean occupancy of true positive binding sites to the mean occupancy of false positive binding sites is 2 (solid) and 10 (dashed) lines. **(E) Variation in auROC with the ratio of mean occupancy between true positive and false positive binding sites. (F) Variation in auROC with sequencing depth**. The ratio of the mean occupancy of true positive binding sites to the mean occupancy of false positive binding sites is set at 2 (solid line) and 10 (dashed line). In (C)-(F), the error bars are the standard deviation computed from ten replicates.

Figures 3C and 3D show the impact of heterogeneity in extraction efficiency and PCR efficiency, respectively, on the auROC of ChIP-seq read count ratios. We found that increases in the mean PCR efficiency did not appreciably increase the auROC. Similarly, an increase in the heterogeneity of PCR efficiency, in terms of its coefficient of variation across the genome, did not decrease the auROC. In contrast, an increase in the mean extraction efficiency, and a decrease in its heterogeneity, led to a higher auROC. Thus, much like in the case of ChIP-seq fidelity, extraction efficiency has a greater impact on the auROC of ChIP-seq read count ratios than PCR efficiency.

Figure 3E shows that as expected, the auROC increases with the ratio of the mean occupancy of true and false positive binding sites. We illustrate in Methods that this mean occupancy ratio can be considered as a proxy for the specificity of the ChIP antibody used, where a higher mean occupancy ratio translates to a higher antibody specificity. A mean occupancy ratio of ~ 10 gives an auROC of around 0.9, which translates to a true positive rate of 0.8 at a false positive rate of 0.07. For a fixed mean occupancy ratio, an increase in the sequencing depth of both ChIP and input samples leads to an increase in the auROC (Figure 3F).

We further evaluated the impact of variation in chromatin accessibility across true and false positive binding sites. Our model of TF-DNA binding in the presence of chromatin assumes that TFs occupy a genomic location only when chromatin is accessible, i.e., the probability of occupancy depends on the energy of the binding site present and the probability that chromatin is accessible at that location (Equation (8)). We found that chromatin accessibility had a low impact on the auROC when the mean accessibility was at least 0.2 (Supplementary Figure 6). We note that equation (8) implicitly assumes that the probability of occupancy at a genomic location is zero in cells where chromatin is inaccessible. This does not hold for TFs such as pioneer TFs [32], which are capable of binding inaccessible chromatin and opening it up for further binding by other TFs. For these TFs, our model under-estimates the occupancy probability at a given genomic location, with the result that the auROC that we obtain in our simulation represents a lower bound.

### Indirect binding affects motif inference and lowers sensitivity more than cooperative binding

The target TF (say, A) of a ChIP-seq experiment is said to be indirectly bound to DNA if it does not bind DNA directly but instead binds a second TF (say, B) that in turn directly binds DNA. Thus, an indirectly bound locus lacks a binding site for the target TF, and the number of reads mapped to this location do not depend on the binding energy of A. This can be seen in the top panel of Figure 4A, where the read count ratios of locations directly bound by A depend on the binding energy of A (*R*^2^ = − 0.73, *p* < 10^-115^) whereas the read counts from indirectly bound regions do not (*R*^2^ = 0.004, *p* = 0.933).

**Figure 4:**
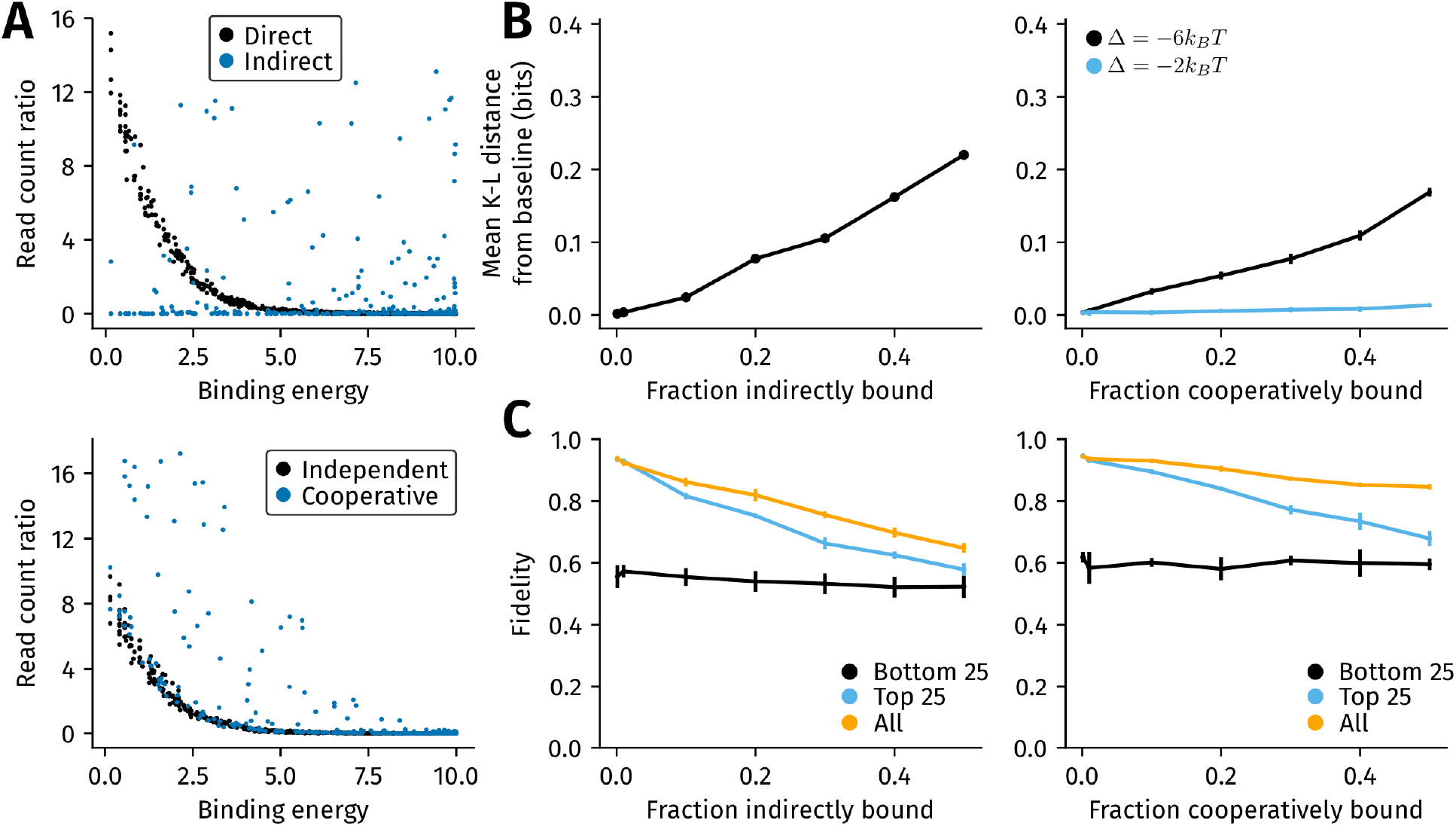
The relative impact of the inclusion of indirectly bound and cooperatively bound locations on motif inference and ChIP-seq fidelity. **(A)** Indirect binding and cooperative binding between TFs A and B are simulated as described in Methods. The scatter plots show the read counts from simulations where 30% of genomic locations are indirectly bound (top panel) or cooperatively bound (bottom panel, interaction energy (Δ) set to −6*k_B_T*). In the top panel, the binding energy of A shown on the x-axis refers to the energy with which A would bind a location if it were in direct contact with DNA. The binding energies of A and B are sampled from a power law (with exponent 0.5) over a range of [0, 10*k_B_T*]. **(B) Inclusion of indirectly bound locations [left] distorts the inferred motif more than the inclusion of cooperatively bound locations [right panel]**. The change in the average K-L distance per base between the baseline motif and inferred motif (y-axis) is plotted against the fraction of indirectly bound regions [left] or cooperatively bound regions [right] on the x-axis. In the right panel, the cooperative interaction energy with the second TF is varied between −6*k_B_T* (black) and −2*k_B_T* (blue) (orange). The error bars are the standard deviation obtained from ten replicates of simulation. **(C) Indirect binding has a greater impact on ChIP-seq fidelity than cooperative binding**. The variation with fidelity and the fraction of indirectly bound [left] or cooperatively bound [right] locations is shown. The fidelity is calculated across all genomic locations (black), the top 25th percentile (blue) and the bottom 25th percentile (orange) of binding sites. The error bars are the standard deviation obtained from ten replicates of simulation.

In contrast to indirect binding, genomic locations that are cooperatively bound by the target TF and the second TF contain a binding site for each TF. The magnitude of the cooperative effect is represented by the interaction energy (Δ, in units of *k_B_T*). Lower (negative) values of Δ represent a larger cooperative effect, with Δ = 0 representing independent binding (see equation (4)). In the bottom panel of Figure 4A, read count ratios of some cooperatively bound locations (Δ = −6*k_B_T*) are higher than independently bound regions. Unlike in the case of indirect binding, however, read count ratios from cooperatively bound regions depend on the binding energy of the target TF *(*R*^2^* = −0.66, *p* < 10^-38^), in addition to depending on the binding energy of the second TF and the interaction energy between both TFs.

In Figure 4B, we separately evaluated the impact of indirect and cooperative binding on the inferred PWM of the target TF. A detailed description of PWM inference in the presence of indirect and cooperative binding is given in Methods. The baseline PWM of the TF Tye7 is first computed with direct binding alone. We then computed the mean K-L distance between this baseline PWM and the PWM inferred in the presence of either cooperative binding or indirect binding with a second TF. We found that the inclusion of a given fraction of indirectly bound sequences distorted the inferred PWM more than the inclusion of the same fraction of cooperatively bound sequences. The divergence of the inferred PWM from the baseline depended both on the binding energy range of the indirectly or cooperatively binding second TF, as well as the strength of the cooperative effect.

Similarly, we found that a given fraction of indirectly bound locations had a greater impact on ChIP-seq fidelity than the same fraction of cooperatively bound locations (Figure 4C). This was true even though the cooperativity is high (Δ = −6*k_B_T*). Indirect and cooperative binding particularly affected the fidelity of read count ratios from the strongest binding sites (the top 25th percentile of read count ratios).

### More than two ChIP-seq replicates are needed to reliably estimate binding energies of low affinity sites from read counts

Next, we assessed the accuracy with which the binding energy at a locus can be estimated using ChIP-seq read count ratios, and the impact of replicates on this accuracy. If we denote the true binding energy of the i-th locus as *ϵ*^(*i*)^ and the estimate of the binding energy as 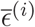, we expect 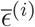 to move closer to *ϵ*^(*i*)^, on average, as more replicates of ChIP-seq are performed. The first step to quantifying the accuracy of 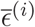 is to compute its posterior probability after having observed read count ratios *R*^(1)^, *R*^(2)^, …, *R*^(*n*)^ from *n* biological replicates, which is the conditional probability 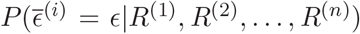. We define the absolute uncertainty or uncertainty of the binding energy estimate to be the 95% Bayesian credible interval of this posterior distribution [33], which we take to be the difference between its 97.5th and 2.5th quantiles. The posterior probability, and hence, the uncertainty of 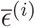 after *n* + 1 replicates are performed, can be computed from 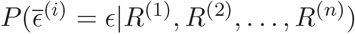 using Bayes’ rule, as shown in Methods.

The expected decrease in the uncertainty of 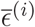 as more replicates are performed can be seen in Figure 5A, where the area under posterior probability of 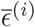 becomes more concentrated around *ϵ*^(*i*)^ with each additional replicate. We repeated this procedure to calculate the reduction in uncertainty at locations with binding energies between 2*k_B_T* and 6*k_B_T* in Figure 5B, and found that the uncertainty is high at sites with the lowest binding energies even after five replicates are performed. At the lowest binding energies, we also checked if the uncertainty after five replicates sufficed to distinguish between two binding sites that differ by one base pair from each other. From an analysis of binding energy matrices of 368 TFs in the BEEML database [13], we estimated that 75% of single mutations change the binding energy of a site by at least 0.24*k_B_T*, while 50% of single mutations change the energy by at least 0.71*k_B_T*. Thus, even three replicates of ChIP-seq would allow us to distinguish between two binding sites that are a single mutation apart with a probability between 50% and 75% at only the relatively high energy binding sites.

**Figure 5:**
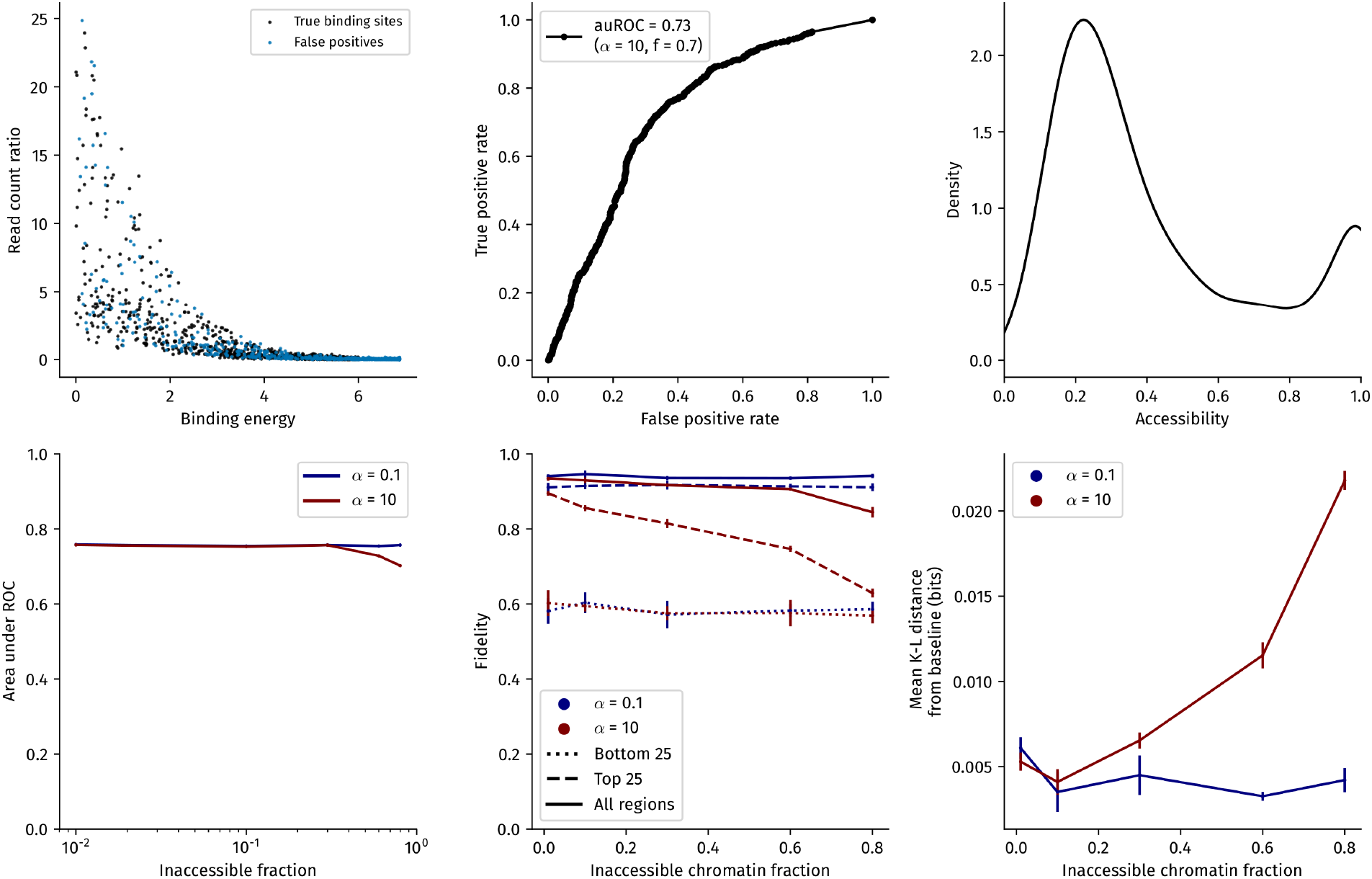
Impact of multiple biological replicates on errors in binding energy estimates. **(A) Posterior estimate of the binding energy at a single genomic locus after 1-4 replicates**. The true binding energy of the locus is *2kBT*, indicated by the vertical dashed line. Four replicates of ChIP-seq were simulated, with the posterior density after each replicate, with the prior distribution set to be the default binding energy distribution employed in all our simulations (Methods). The blue lines indicate the absolute uncertainty of the binding energy estimate, which we define as the 95% Bayesian credible interval of the binding energy. This is calculated as the difference between the 2.5th and 97.5th quantiles of the posterior density. **(B) The reduction in absolute uncertainty of posterior binding energy estimates of five locations, each with a different binding energy, after three replicates of ChIP-seq**. The error bars are the standard deviation in estimates of the absolute uncertainty calculated after 100 trials; the filled circles represent the mean absolute uncertainty from these trials. The dashed red lines represent the values of absolute uncertainty required to distinguish between binding sites sequences that are a single mutation apart at least 50% of the time (upper,0.71*k_B_T*) and at least 75% of the time (lower,0.24*k_B_T*).

### Simulating FASTQ reads using ChIPulate

In addition to simulating read counts from ChIP and control samples as described so far, ChIPulate can output single-end and paired-end reads for each genomic location if its chromosome coordinates are specified.

Figure 6A illustrates the working of ChIPulate when it is run in FASTQ generation mode. To generate sequence reads, a set of *n* genomic intervals (*b*_1_, *e*_1_), (*b*_2_, *e*_2_), …, (*b_n_*, *e_n_*), along with a set of “summits” *s*_1_, *s*_2_, … *s_n_* is input to ChIPulate. These summits can be thought of as the physical location of TF-DNA binding in each interval. The total and unique read counts (*r_i_* and *u_i_*, respectively) at each interval are computed as described in Methods based on the binding energy (or energies) associated with each interval. Fragments that are PCR duplicates are tracked and the reads arising from duplicate fragments are named appropriately to reflect their status as a PCR duplicate. In the ChIP sample, the start positions of fragments in the i-th interval are sampled from a Gaussian distribution with mean *s_i_* −*d*/2 and standard deviation *j*, where *d* is the fragment length and *j* is referred to as the fragment jitter. This captures the notion that a genomic position close to a TF-DNA binding event is more likely to give rise to a fragment in the ChIP sample than a genomic position that is far away. In the control sample, each position in an interval is assumed to be equally likely to give rise to a fragment. This corresponds to the Poisson background model assumed by peak callers such as MACS2 [24]. Thus, in the control sample, ChIPulate samples start positions in the i-th genomic interval from a Uniform(*b_i_*, *e_i_*) distribution. The *l* base pairs atthe ends of each fragment in both ChIP and control samples represents the read length. Though Figure 6A illustrates the simulation of paired-end reads, ChIPulate can simulate both single-end reads and paired-end libraries, where the sequence of each read is assumed to be error-free.

**Figure 6:**
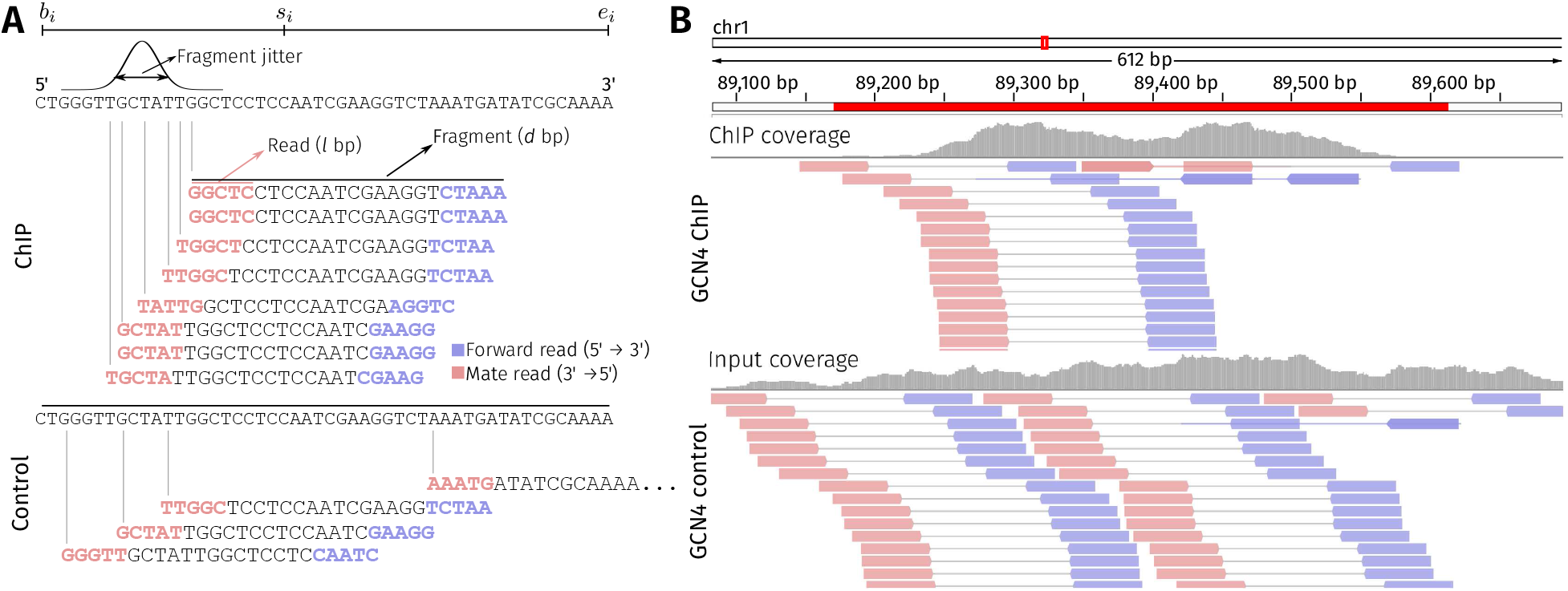
Example of simulating sequence reads using ChIPulate. **(A) Illustration of ChIPulate’s read generation method**. The i-th genomic location is a genomic interval (*b_i_*, *e_i_*), with the “summit” of the interval located at s_i_. The total read count (*r_i_*) and unique read count (*u_i_*) at the i-th interval in the ChIP and control samples are calculated as described in Figure 1. The starting positions of fragments in the ChIP sample are drawn from a Gaussian distribution with mean *s_i_* − *d*/2 and standard deviation *j*, where *d* is the fragment length and *j* is the fragment jitter. In the control sample, fragment start positions are drawn from a Uniform[*b_i_*, *e_i_*] distribution. The Phred-33 quality value of every base is set to be 75, which corresponds to a base-calling error probability of 10^-42^. The read length is set to *l* bp, and both paired and single end reads can be simulated. **(B) Paired-end reads simulated from genomic intervals containing an experimentally determined GCN4 binding site**. The genomic intervals containing GCN4 binding sites were taken from an earlier publication [34]. Paired-end reads of 50 bp in length were simulated using ChIPulate, with the fragment jitter set at 50 bp and the fragment length set at 200 bp. The binding energy of the highest affinity GCN4 binding site in each interval was computed using the GCN4 binding energy matrix from the BEEML database. The remaining ChIPulate parameters were set at their default values.

Figure 6B is an IGV (Integrated Genomics Viewer) [35] snapshot of paired-end reads simulated from a genomic interval containing a GCN4 binding site in the *S. cerevisiae* genome. In this example, we chose a set of 549 genomic regions that were found to be bound by GCN4 in an earlier publication [34], and computed the binding energy of GCN4 in each region using its binding energy matrix from the BEEML database. The starting position of the lowest energy binding site was set to be the summit of each region. We then generated the start positions of fragments (200 bp in length) in the ChIP sample by setting the fragment jitter to 50 bp. The read length for the paired-end reads were set at 50 bp at a sequencing depth of 100. The sequence of each read was assigned based on the sequence of the S288C R64-2-1 reference of the *S. cerevisiae* genome [36] from the *Saccharomyces* Genome Database [37]. The reads generated were aligned to this genome reference using the BWA aligner [38].

## Discussion

ChIPulate substantially and qualitatively extends previous statistical models of read counts obtained in a ChIP-seq experiment. Early work by Zhang et. al assumed a distribution of binding affinities across the genome and computed occupancy using a similar biophysical model to ours [24], but did not simulate extraction and amplification. Bao et. al explicitly accounted for the effect of extraction efficiency on read counts [26], but did not model the underlying biophysical occupancy of genomic loci based on their binding energies. More recent work by Ruan et. al, accounted for the effect of chemical potential on TF-DNA occupancy and its impact on motif inference [14]. However, the authors evaluated the impact of experimental noise by adding Gaussian noise to the energies of binding sites rather than modeling the downstream processes of the ChIP-seq protocol.

Our pipeline combines a biophysical model of TF-DNA binding with a detailed simulation of fragment extraction, amplification and sequencing. This allows us to analyze the impact of heterogeneity in fragment extraction and PCR amplification on the motif inferred from bound genomic loci, and the fidelity of read count ratios in discriminating relatively close binding energies. We also evaluated how these factors, along with chromatin accessibility and sequencing depth, affect the false positive peak detection rate. The role of biological factors such as indirect binding and cooperative binding, which are frequent occurrences in ChIP-seq datasets, were also evaluated for their impact on motif inference and fidelity. Finally, we measured the accuracy of using read count ratios to infer the energy of a binding site and calculated the improvement in this accuracy after several biological replicates are performed.

In our simulations, fragment extraction encapsulates multiple steps in the ChIP-seq protocol, namely, cross-linking TF-DNA bound complex, antibody-mediated pull down of fragments bound by the TF, and the removal of cross-links. In addition to the mean efficiency, the nature of the extraction heterogeneity dictates the magnitude of its impact on motif inference and fidelity, with power law distributions having a greater impact than normally distributed heterogeneity. The effect of extraction heterogeneity on motif inference also depended on the structural class of the TF, where we found that a low mean extraction efficiency distorted motifs with a lower information content much more than those with high information content. However, for a given TF, it is unclear as to whether a normal or power law distribution better capture experimental variation in extraction efficiency across the genome. Previous reports suggest that variation in chromatin accessibility alone imposes a power-law distribution on extraction efficiency variation [18]. The development of alternate protocols that reduce the number of extraction steps [39, 40, 41] would improve the fidelity of read count ratios and reduce the number of false positive peak calls. In contrast to extraction efficiency, variation in PCR amplification efficiency had a much lower impact on motif inference and the fidelity of read count ratios. However, in the presence of sequencing errors during amplification (which our model does not currently include) and imperfect mappability, a large mean amplification ratio in practice would increase false positive peak calls [42] and likely affects fidelity more than indicated by our simulations.

Though we explicitly modelled TF-DNA binding here, our conclusion on the impact of extraction and amplification efficiency on fidelity also likely holds for the ChIP-seq of histone marks. Though the occupancy of histones in nucleosomal complexes would have to be modelled in a very different manner to that of TFs [43, 44, 45, 46], this only affects the calculation of 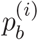. The remaining steps of extraction, PCR amplification and sequencing that are carried out in the ChIP-seq of a histone mark are identical to those that are simulated here. An issue of interest in a histone ChIP-seq dataset is the method employed to process the control sample [47, 48], where employing an antibody against the histone H3 in the control sample provides a better estimate of *p_ext_* than an input sample [47]. Our simulations suggest that this would be true only if the mean extraction efficiency of fragments with these methods is not too much lower than using an input sample. This is because a low mean value of *p_ext_* in the control sample would poorly sample any potential variation in extraction between genomic regions in the ChIP sample.

Extraction of DNA fragments due to indirect binding is believed to be frequent in ChIP-seq experiments [49, 10]. We find that the inclusion of indirectly bound regions has a more adverse effect on motif inference and fidelity than even strongly cooperatively bound regions. The extent of indirect binding in a dataset depends both on antibody quality [23, 11] and the cross-linking protocol employed [50] except when the ChIP-seq target protein is a co-factor which, by definition, is always indirectly bound. When antibody quality leads to indirect binding, we note that indirectly bound peaks and false positive peaks are hard to distinguish. Our work suggests that minimizing the extent of indirect binding and developing computational methods to detect them may provide a potent avenue to improving the fidelity of ChIP-seq experiments. For instance, protocols such as ChIP-exo [51, 11, 52] allow indirectly bound regions to be more easily detected through the peak shapes obtained after alignment and represent an important direction for further improvement of the ChIP-seq protocol.

Our results shows that several replicates of ChIP-seq are required if the energy of a binding site is to be inferred through read count ratios within a certain confidence interval. Since our simulation assumes a single site per peak, it should be noted that a single replicate of ChIP-seq can suffice to accurately infer a PWM for the TF [53], which in turn reliably estimates the energy of the binding site [54, 55, 56]. However, in the more general case where a peak may harbor multiple TF binding sites, the mapping between the binding energies of sites and the read count ratios in such a region is far more complex than in our simulation. This means that reliably inferring the occupancy at such a locus through read count ratios is likely to involve more replicates than we have estimated here. Since our simulations in this calculation assumed relatively low heterogeneity in extraction and PCR efficiency, further improvement in these steps of the protocol may not be the most effective avenue to reduce the number of replicates required to infer occupancy. Instead, reduction in sequencing cost to obtain multiple biological replicates of ChIP-seq will yield an accurate inference of *in vivo* occupancy at a genomic locus.

There is considerable scope for extending ChIPulate. We assumed that amplified fragments can be de-duplicated after sequencing in our simulation, which is possible with paired-end sequencing and unique molecular identifier (UMI) based methods of PCR amplification [57, 58, 59]. Most ChIP-seq libraries are, however, subjected to single-end sequencing. It might be of interest to investigate the effect of imperfect de-duplication with single read sequencing. Furthermore, we did not take into account differences in mappability across the genome or sequencing errors, which can cause reads originating from a genomic locus to not map back to the same region [60]. These are relatively easy to account for in our simulation since it keeps track of individual fragments when they are extracted, amplified and sequenced. Finally, the FASTQ generation mode of ChIPulate can be used to comprehensively benchmark ChIP-seq peak callers and other computational pipelines that process ChIP-seq data.

Overall we have presented a detailed ChIP-seq simulation model and software pipeline that can be extended in various ways, including other bulk and single-cell sequencing protocols.

## Acknowledgements

We thank Gautam Menon, Sandeep Krishna, Parul Singh and Aswin Sai Narain Seshasayee, for discussions. SH was funded in part by NSF award 1564785. VD was supported by the Simons Foundation. RS was supported by the PRISM 12th plan project at the Institute of Mathematical Sciences, funded by Department of Atomic Energy, Government of India.

## Methods

### ChIP-seq simulation framework Genome-wide TF-DNA binding model

In the most basic version of our simulation, we consider the genome to be a set of *n* locations. Each of these locations contains a single binding site for the ChIP-seq target TF. We follow an approach similar to [27], where, at a given point in time, the i-th locus can be in one of two states — bound by the target TF or unbound. We assign a binding energy *ϵ_u_* to the unbound state and a binding energy *ϵ*^(*i*)^ to the bound state. The Boltzmann weights of both these states are then exp(−*ϵ_u_*/*k_B_T*) and exp(−(*ϵ*^(*i*)^ − *μ*)/*k_B_T*), respectively. *k_B_* is the Boltzmann constant *T* is the temperature at which the binding occurs, and *μ* is the chemical potential of the TF, which is proportional to the logarithm of the concentration of the TF [27]. The probability of finding the i-th locus in the bound state is then

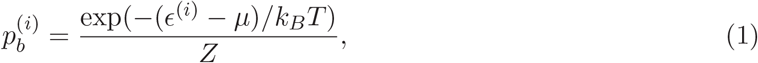

where *Z* = exp(−(*ϵ*^(*i*)^ − *μ*)/*k_B_T*) + exp(−*ϵ_u_*/*k_B_T*). This can be re-written as

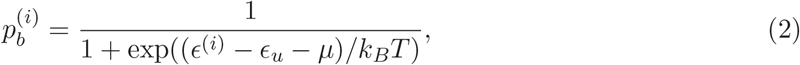

Thus, the occupancy at a location is determined by the three quantities *ϵ*^(*i*)^, *ϵ_u_*, *μ*. In the convention followed in this paper, the scale of *ϵ*^(*i*)^ is chosen such that the highest affinity binding site has a binding energy of 0, with more positive values representing weaker binding sites. We set *μ* = 3*k_B_T*, which is within the range of values suggested by an earlier calculation in [27]. Finally, we set *ϵ_u_* = 1.59*k_B_T*, which ensures that at the location with the highest affinity site i.e., when *ϵ*^(*i*)^ = 0, the occupancy probability 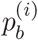 is 0.99.

*ϵ*^(*i*)^ can be thought of as a mismatch binding energy, where mutations that change the binding site sequence away from the highest affinity sequence increase its energy and thus lower the probability that it is occupied. This is in line with the convention followed in the binding energy matrices in the BEEML database.

To compute the probabilities of binding at all *n* locations, 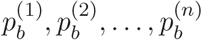, using equation (1), we assign a binding energy to each locus. We sample binding energies from a power law that is truncated to a specified range. We define the probability density function f of this truncated power law, with parameters 1 ¿ *α* > 0, *E_max_* > 0, as

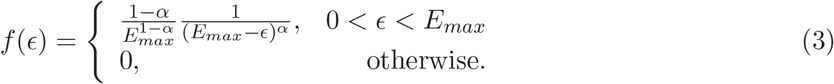

Our choice of such a power law is motivated by the analysis of binding energies of sites that are occupied by the TFs CRP and FNR in the *E. coli* genome [61]. Unless stated otherwise, we set *α* = 0.5 and *E_max_* = 10*k_B_T*. This corresponds to a situation where most of the binding sites in the genome have a low affinity (or high energy) for the target TF. *E_max_*= 10*k_B_T* typically corresponds to a binding site that is 3-4 mutations away from the strongest binding site, since each mutation to the strongest sequence typically adds an energy 1 − 3*k_B_T* to it for most TFs [27]. We set the chemical potential of the target TF, *μ*, to be 3*k_B_T* by default in all our simulations in the main text.

In a population of *C* cells, when the i-th locus has a probability 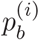 of being bound, then the number of bound fragments extracted from the i-th locus, denoted 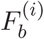, follows a Binomial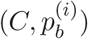 distribution, with the mean number of bound fragments is 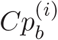.

#### Input sample

In most ChIP-seq experiments, a small number of cells are used to carry out a control, or input, experiment where fragments from cells are extracted without the use of antibody specific to the target TF. The read counts obtained in the input experiment are used to normalize, in a region or locus-specific manner, the read counts obtained from the ChIP sample. The input sample helps take into account heterogeneity that may arise in the extraction, amplification and sequencing of fragments from different genomic locations. The difference between the input and ChIP samples is in the number of cells employed and the probability of occupancy employed for each location. We assume that 10% of the number of cells used in the ChIP experiment are used in the input sample. The occupancy of genomic regions in the background sample consists of the nonspecific binding of all the TFs in the cell, along with different sets of TFs that may be specifically bound at each genomic location. We phenomenologically model this scenario by assigning a fixedbackground binding energy 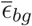 to each of the *n* genomic locations, from which we use equation (1) to compute the probability of occupancy 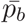. We set *μ* = 0 in equation (1) for the input sample and keep *ϵ_u_* = 1.59*k_B_T*.

If the number of cells used in the input sample is *C′*, then the number of bound fragments at each location follows a Binomial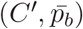 distribution. Unless stated otherwise, we set 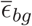 to 1*k_B_T* in simulations where we do not model false positive binding sites in the ChIP sample. When we simulate false positive binding sites, we set 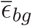 to a value where the ratio between the mean probability of occupancy in the ChIP sample to the mean probability of occupancy in the input sample is 10.

Aside from these differences in the TF-DNA binding model in the input and ChIP samples, the simulation procedures used in the description of the extraction, amplification and sequencing steps are common to both samples.

### Cooperative binding

Consider TFs A and B, where A is the target of the ChIP-seq experiment, however, its binding depends on nearby binding of TF B. To simulate such cooperative binding between A and B, we have each of the *n* genomic locations in our simulation to contain a single binding site each for A and B, with binding energies 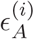 and 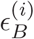 and an interaction energy Δ between them.

A given genomic locus can be in one of four states — unbound, bound by A, bound by B, or bound by both A and B. The probabilities of the locus being in each of these four states is [62]

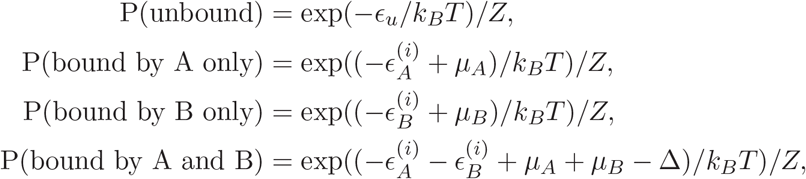

where 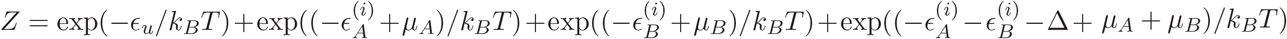. Δ = 0 corresponds to independent binding between A and B, Δ < 0 corresponds to cooperative binding between A and B and Δ > 0 represents competitive binding between A and B.

Since the ChIP experiment involves extracting fragments that are bound by A, the probability of a location being bound by A is the sum of the probability of the location being bound only by A and the probability of being bound by A and B. Thus, the occupancy probability of the i-th location, 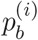, as seen in the ChIP-seq of A is :

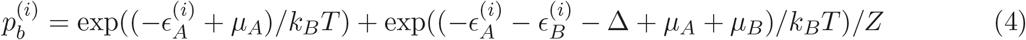

We assume here that the binding site of A is not occupied by B, and vice versa. Further, we assume that there is no cooperativity between two distinct genomic locations *i* and *j*. We do not take the chemical potential of individual TFs into account, and thus implicitly assume that the concentrations of both TFs are similar.

### Indirect binding

Consider TFs A and B, where A is the target of the ChIP-seq experiment. We say that A indirectly binds DNA if A binds to B which in turn binds DNA. To simulate indirect binding, we assume that the probability of occupancy 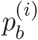 is proportional to the binding energy of B at the i-th location, 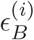 as

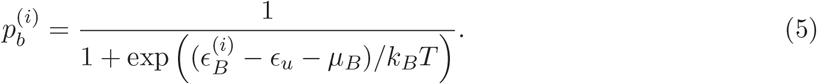

On the other hand, at locations directly bound by A, 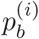 is dependent on 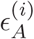 according to equation (1). We set the chemical potential of A and B to zero and assume that B cannot occupy binding sites of A, and *vice versa*.

### False positive binding sites

We consider a detected binding site to be false positive if it is bound by a TF (or TFs) other than the target TF of a ChIP-seq, or not bound by any TF at all, but fragments from which are nonetheless extracted in Step 2 of simulation. This differs from indirect binding, where the target TF is part of the complex of TFs that is bound to DNA. False positives reflect antibody quality or specificity.

In our simulations of false positive binding sites in Figure 3, we set the number of false positive binding sites to be equal to the number of true positive binding sites. We note that the area under the ROC curve is independent of the number of false positive binding sites simulated.

We set the binding energy distribution of true positive binding sites to be a power law between 0 and 6*k_B_T* with *α* = 0.5. This differs from the default power law distribution employed for true positive binding energies in our other simulations. This was because we found that binding sites with energies in the range 7 – 10*k_B_T* had read count ratios that were very close to each other. Thus, when we computed the ROC curve with binding sites in this energy range, we saw that the true positive rate showed an abrupt jump as the threshold on the read count ratio was lowered below a point. Such jumps in the true positive rate make it difficult to compare ROC curves. By restricting the energy range of true positive binding sites to between 0 and 6*k_B_T*, we ensured that the true positive rate did not display this jump-like behaviour, which in turn allowed us to use the auROC as an accurate measure of detection performance.

We set the binding energies of the false positive sites to follow a power law distribution where the minimum binding site energy was 0*k_B_T*. The maximum binding energy 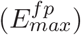 and *α_fp_* was set such that the ratio of the means of true and false positive binding energies could be fixed at a constant value while the variances were kept equal. Suppose the true positive binding energy distribution follows a power law over the energy range 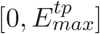 and exponent α_tp_ such that its mean is *m* and variance is *v*. Then, setting *α_fp_* and 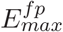 as

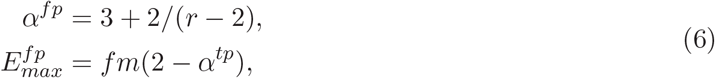

where *r* = *v*/(*g*^2^*m*^2^) + 1, sets the mean false positive binding energy to be *fm* while the variance remains at *v*.

Once the true and false positive binding energies are sampled, we compute the ratio between the mean occupancy of the true positive sites to that of the false positive binding sites. This is calculated as

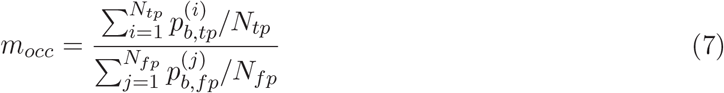

where *N_fp_* and *N_tp_* are the number of false and true positive binding sites, respectively, and 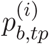 and 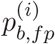 are occupancies at true and false binding sites respectively.

In Figure 3, we set *α_fp_* = 0.76 and 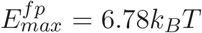, which corresponds to a mean occupancy ratio of ~ 2. This is in line with the antibody recommendations of the ENCODE consortium [23], whose recommendation for antibody quality is that the primary reactive band should be at least 50% of the signal on the blot, which we interpret to represent a mean occupancy ratio between true positive and false positive sites in the ChIP sample to be at least 2.

Finally, we set the background binding energy, which determines the occupancy of fragments in the input sample, to a value of 4.96*k_B_T*. This sets the ratio of the mean occupancy of true positive sites in the ChIP sample to the mean occupancy of the same sites in the input sample to 10. This is in line with the 5– to 13– fold-enrichment observed in read counts of ChIP-seq peaks between the ChIP and input samples in the ENCODE project [23].

### Chromatin accessibility

We modeled chromatin accessibility using a sigmoidal prior based on DNAse-seq hypersensitivity data following [63, 3]. Briefly, the occupancy probability calculated based on the binding energies in equation (1) is multiplied by the sigmoidal function—

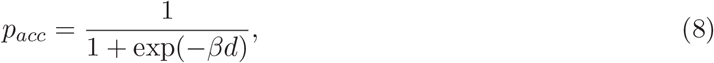

where *d* is the DNAse-seq read count density, which is average per-base DNASe-seq read coverage within a 150 bp window.

Thus, for a genomic locus whose occupancy probability is *p_b_*, the probability that it is bound in Step 1 of our simulation is the product *pbpacc*. Thus, the occupancy probability essentially provides a scaling factor for binding probability.

Instead of simulating a profile of DNAse-seq read counts across the genome, we used read counts from DNAse-seq data of the *M. musculus* DBA/2 cell line (accession number ENCFF871YIT, lab of John Stamatoyannopoulos), available from the ENCODE Consortium [64, 65]. We used the filtered binary alignment map (BAM) files, which were aligned to the mm10 genome assembly using the ENCODE data processing pipeline. We chose the DNAseq-seq profile of this cell line as a reference DNAse-seq profile since it cleared all quality audit error categories laid out by the ENCODE Consortium for DNAse-seq data.

In order to consider only regions with accessible chromatin, we counted the number of reads falling into non-overlapping 100 kb windows of the genome and retained the top 10% of windows that had the highest read counts. We finally counted the reads falling into non-overlapping 150 bp windows and stored these read counts. In each ChIP-seq simulation, we uniformly sampled read counts of *n* of these regions, where *n* is the number of binding locations being simulated, and plugged them into equation (8) (after dividing the counts by 150) to compute chromatin accessibility.

### The extraction process

Given the number of bound fragments 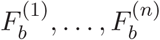, each fragment needs to be extracted from the cell population. This extraction process involves several steps that include the lysing of cells to extract DNA, cross-linking of bound proteins to DNA, the size selection of sheared fragments, etc. After each of these steps, the number of bound fragments either stays the same or reduces. Our model of fragment loss during these steps tracks the reduction in fragment numbers prior to PCR amplification step.

We assume that the number of extracted fragments from the i-th genomic locus, denoted 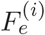, follows a Binomial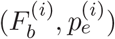 distribution. 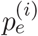 is the probability of a bound fragment from the i-th region being successfully extracted and present in the pre-PCR fragment pool. *p_e_* can vary across the genome [66, 67] and is also dependent on the extraction procedure employed [68]. Unless stated otherwise, we assume that *p_e_* follows a normal distribution truncated to lie between 0 and 1, with a mean of 0.5 and a standard deviation of 0.05.

Since we assume that 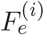 is a binomial sample of 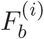 fragments, the addition of more extraction steps into the simulation only changes the value of 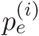 without necessarily violating the binomial assumption. Suppose we add an additional extraction step with efficiency *e_i_* into the simulation such that 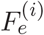 ~ Binomial(*X_i_*, *e_i_*), where *X_i_* ~ Binomial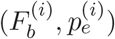. Then, by the laws of total variance and total expectation, the mean of 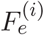, for a fixed value of 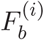, is 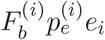 and its variance is 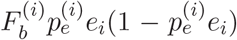. By setting 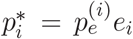, we see that 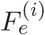 (conditioned on 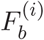) follows a Binomial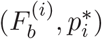 distribution up to the first two moments. Thus, the parameter 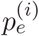 can be thought to implicitly take into account a variety of steps employed in the extraction process.

As stated in the previous section, we employ the same values of 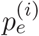 in both ChIP and input samples. In a single run of the simulation, it can happen that the number of fragments extracted in the ChIP sample exceeds that of the input sample, or vice versa. In practice, since fragments are typically more efficiently extracted in the input sample than in the ChIP sample, DNA extracted from the input library is diluted such that the number of fragments in both ChIP and input samples are equalized before amplification.

In our simulation, we compute a down-sampling factor 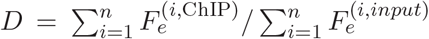, which is the ratio of the total number of extracted fragments in the ChIP sample to that in the input sample. If *D* < 1, we down-sample extracted fragments in the input sample by replacing 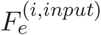 with the sample *X_i_* ~ Binomial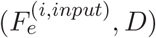. If *D* > 1, we similarly down-sample the fragments from the ChIP sample instead of the input sample.

### Mean occupancy ratio as a measure of antibody specificity

In equation (7), we defined the mean occupancy ratio as the ratio between the mean occupancy of true positive binding sites to that of false positive binding sites. This ratio can also be interpreted as a measure of antibody specificity in the following way.

Consider a genome with *n* true positive binding sites and *m* false positive binding sites. Suppose the extraction efficiency is 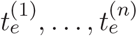 at true positive binding sites and 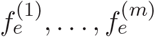 at false positive binding sites. Then, the average number of fragments extracted from the i-th true positive binding site in the ChIP sample is 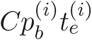, and the average number extracted from the j-th false positive binding site is 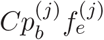, where 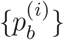 are occupancies at each location. The ratio (*r_e_*) between the average number of fragments extracted from true positive binding sites to that from false positive binding sites is then

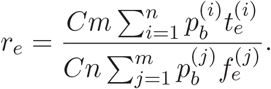

This is the ratio that determines the accuracy of ChIP-seq in distinguishing between true and false positive binding sites. If *r_e_* is large, true positive binding sites will give rise to more fragments than false positive binding sites, leading to true positive sites having larger read count ratios. A high value of *r_e_* is achieved if, for 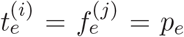 across the genome, true positive sites have a larger mean occupancy than at false positive sites. Setting 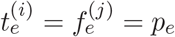 in the above equation reduces *r_e_* to the mean occupancy ratio defined in equation (7).

Conversely, if 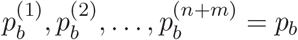 across the genome, then *r_e_* reduces to the ratio

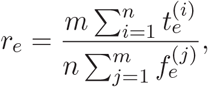

which is merely the ratio between the mean extraction efficiency at true positive binding sites to that at false positive binding sites. This constitutes a measure of the specificity of the antibody employed against the target TF of the ChIP-seq. Thus, ensuring that the average extraction efficiency at true positive sites is higher than at false positive sites ensures that the former give rise to higher read count ratios.

### PCR amplification

We simulate PCR amplification using the model in [69] which we briefly explain below. In this model, each DNA fragment has a probability *p_a_* of being amplified in a cycle of PCR and gives rise to two fragments. The fragment fails to undergo amplification and remains as a single fragment with probability 1 − *p_a_*.

Suppose *S_n_cy__* is a random variable that represents the number of amplified fragments after *n_cy_* cycles of PCR. *S_n_cy__* can assume values between 1 and 2*^n_cy_^*.

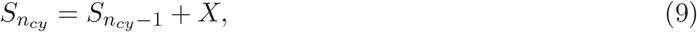

where X ~ Binomial(*S*_*n*_*cy*−1__, *p_a_*). If we represent S_n_cy_−1_ as a vector of length 2^*n*_*cy*−1_^, the probability distribution of *S_n_cy__* can then be calculated from *S*_*n_cy_*−1_ in terms of an updating matrix *M*^(*n_cy_*)^ (of dimension 2^*n_cy_*^ × 2^*n_cy_*−1^) by setting *S_n_cy__* = *M*^(*n_cy_*)^*S*_*n_cy_*−1_ where

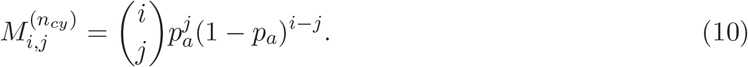

Thus, for a given value of *p_a_*, the distribution of *S_n_cy__* can be computed recursively from *S*_*n*_*cy*−1__, *S*_*n*_*cy*−2__, …, *S*_1_ when we start from a single DNA fragment. For computational efficiency, for values of *n_cy_* between 10 and 15, we precomputed and stored the distributions of *S_n_cy__* for values of *p_a_* between 0.01 and 0.99 in steps of 0.01. In order to simulate *n_cy_* cycles of PCR amplification for 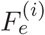 fragments, we draw 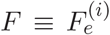 samples 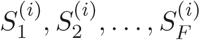 from the distribution *S*_*n_cy_*_ corresponding to the value of 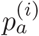 at that locus.

The mean and variance in the number of amplified fragments, 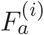, starting with a single fragment at the first cycle and an amplification efficiency of *p_a_*, can be calculated from branching process theory [70]. After a single cycle of PCR, the mean (*m*_1_) and variance (*v*_1_) are

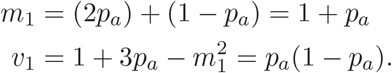

After *n_cy_* cycles of amplification, *m_n_cy__* and *v_n_cy__* are given by —

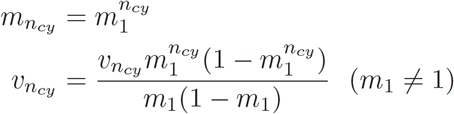

Substituting the expressions for *m*_1_ and *v*_1_ into the above equation, we have

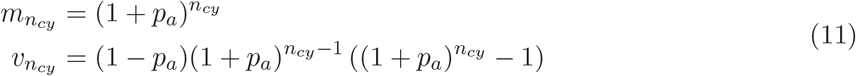

We refer to *A* ≡ *m_n_cy__* = (1 + *p_a_*)*^n_cy_^* as amplification ratio of PCR at that location.

We assume that the PCR efficiencies at a locus remains constant across all cycles of amplification. We also assume that the number of amplified fragments obtained from each starting DNA fragment is statistically independent of amplifications at other fragments. This implies that starting with 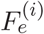 fragments, the mean number of amplified fragments after *n_cy_* cycles of PCR is 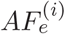. The expressions in equation (11) are valid only when PCR has not yet reached a saturation of amplification where the concentration of PCR primers, polymerase, etc. relative to the concentration of the amplified fragments is enough to ensure that *p_a_* does not decrease as *n_cy_* is increased.

By default, we set *A* = 1000 and *n_cy_* = 15 across the genome in all our simulations unless stated otherwise. This corresponds to a value of *p_a_* that is ≈ 0.58.

### Sequencing

At the end of the PCR amplification step, there are 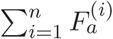 amplified fragments from all *n* genomic locations. In this set of amplified fragments, there are 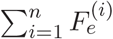 unique fragments, each of which come from different cells, and the remaining 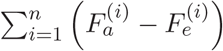 fragments are duplicates obtained through PCR. During the sequencing step, a total of *r = cn* fragments are sampled from this amplified pool and sequenced. *r* denotes the total read count of the experiment and *c* is the sequencing depth.

Since each fragment has an equal probability of being sampled, the read count sample *r*_1_, *r*_2_, …, *r_n_* from amplified fragment pools 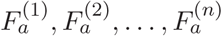, where 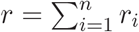, follows a multivariate hypergeometric distribution. To draw a sample from this distribution, we implemented the following procedure [71]—

Given two pools of fragments of pool sizes *A* and *B* from which r reads are to be sampled, the probability that k reads come from the first pool is

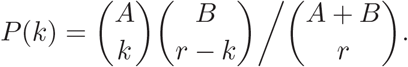

We extend this distribution to sampling from *n* pools of fragments, each of size 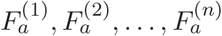 by first drawing *r* reads from two pools of size 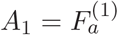 and 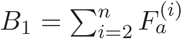. The probability of observing r_1_ reads from the first locus (i.e. pool *A*_1_) is

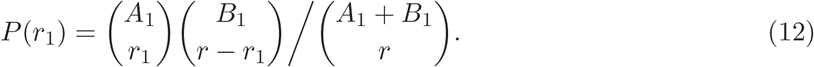

To draw reads for the second locus, we draw a sample of size *r − r*_1_ from two pools of size 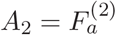 and 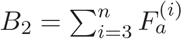.

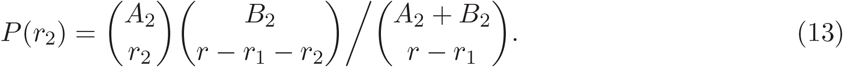

We continue this sampling process till all *r* reads have been sampled.

The read count at the i-th location, *r_i_*, consists of a mixture of *u_i_* unique reads and *d_i_* duplicate reads such that *r_i_* = *u_i_* + *d_i_*. We keep track of the fragments that were obtained via amplification versus those obtained in extraction step, and therefore, by simulation design, the *u_i_* reads can be perfectly separated out from the *d_i_* reads, which is detailed below.

As stated in the earlier section, 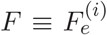 fragments extracted from the i-th location undergo PCR amplification to give 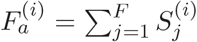 fragments, where 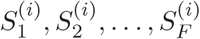 are the number of amplified fragments obtained from each of the 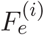 fragments. To sample *u_i_* we first draw samples 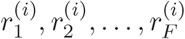 from the fragments pools 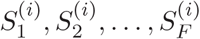 such that 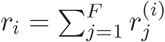. We can write 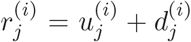, where 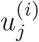 and 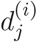 are the numbers of unique and duplicate fragments sampled from 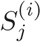. Since we assume that each of the 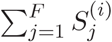 fragments has an equal probability of being sampled, 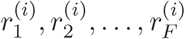 is a multivariate hyper-geometric sample of 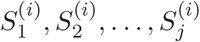, which we generate using the same procedure outlined for sampling *r*_1_, *r*_2_, …, *r_n_* reads from 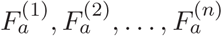 fragments.

We then compute *u_i_* for each location as follows. If 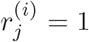, then 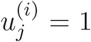 and 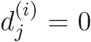. Since only one fragment in 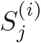 is unique, 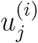 cannot exceed 1, which means that if 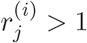, then, 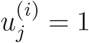 and 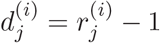. We can then obtain *u_i_* by setting 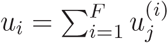.

In the process of simulating read counts, We do not account for mutations than can cause alignment errors in actual ChIP-seq reads, which in turn can change read counts at a genomic locus.

### FASTQ read generation

To generate sequence reads, a set of genomic *n* intervals (*b*_1_, *e*_1_), (*b*_2_, *e*_2_),…, (*b_n_*, *e_n_*), along with a set of “summits” *s*_1_, *s*_2_,…, *s_n_* need to be input to ChIPulate. These summits can be thought of as the physical location of TF-DNA binding in each interval. The total and unique read counts (*r_i_* and *u_i_*, respectively) at the i-th interval are computed as described above, based on the binding energy (or energies) associated with each interval.

In the ChIP sample, after *r_i_* and *u_i_* are computed, *u_i_* fragment start positions 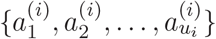 are sampled from a Gaussian distribution with mean *s_i_*– *d*/2 and standard deviation *j*, where *d* is the fragment length and *j* is referred to as the fragment jitter. Suppose the copy number of each of the *u_i_* fragments is *c*_1_, *c*_2_,…, *c_u_i__*, such that *c*_1_+ *c*_2_+…+ *c_u_i__* = *r_i_*, where *c_i_* ≥ 1. Then, a set of *r_i_* fragment start positions can be obtained by repeating the j-th unique fragment start position *c_j_* times to give the set (*a*_1_,…, *a_c_1__*, *a*_2_,… *a*_c_2__,…, *a_u_i__*,…, *a_c_i__*}. The fragment jitter parameter *j* controls the dispersion of the fragment start positions from the position *s_i_*.

The fragment start positions are simulated differently in the control sample. Here, given a unique read count *u_i_* and total read count *r_i_* at the i-th genomic interval, unique fragment start positions (*a*_1_, *a*_2_,…, *a_u_i__*} are sampled from a Uniform(*b_i_, e_i_*) distribution. Given this set of unique fragment positions, the set of *r_i_* fragment positions is constructed in the same way as in the ChIP sample, with the j-th unique fragment’s starting position *a_j_* being repeated *c_j_* times.

Suppose the read length being simulated is *l* base pairs. At the end of the previous step, we have a total of *r* = *cn* fragment start positions each in the ChIP and control samples, where *c* is the sequencing depth and *r* is the total read count. Suppose the the start position of the i-th read is *a_i_* and the genome is a *N* letter sequence *g*_1_*g*_2_…*g_N_*, with the reverse complement sequence denoted 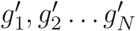. Then, if single-end sequencing is being simulated, we assign the sequence read to begin from the positive strand or the negative strand with equal probability. If the strand assigned to the read is positive, we assign the sequence *g_a_i__g_a_i_+l_*… *g_a_i_+l_* to the i-th read. If the strand assigned to the read is negative, we assign the sequence 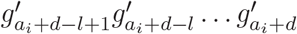. If paired-end sequencing is being simulated, we generate r read pairs, i.e., 2*r* reads in total, where each fragment gives rise to one read from the positive strand with the sequence *g_a_i__g_a_i_+1_* … *g_a_i_+l_* and one read from the negative strand with the sequence 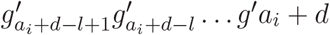. ChIPulate uses the `getfasta` subcommand of `pybedtools` from the input genome FASTA file to extract these sequences [72, 73].

The quality of each base call in every sequence read is set to be K, the maximum possible value on the Illumina Phred-33 quality scale. This corresponds to a base call error probability of 10^-42^ [74].

### Default values

Unless stated otherwise, we use the following values by default—

#### Genome-wide TF-DNA binding model

*n* = 1000, *ϵ*_1_,…, *ϵ_n_* are sampled from a power law (with *α*= 0.5) between 0 and 10*k_B_T*, 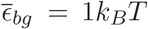, *C* = 10^5^(ChIP sample), *μ*= 3*k_B_T, C* = 10^4^ (input sample)

#### Extraction process

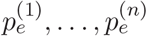 are sampled from a *N*(0.5, 0.025) distribution that is truncated to lie in [0, 1]. Further, at each location, the extraction efficiency is set to be the same in both ChIP and input samples.

#### Amplification process

*A* = 1000, *n_cy_* = 15

#### Sequencing

*c* = 100

### Motif estimation

To simulate motif estimation, we first sample binding energies *ϵ*_1_, *ϵ*_2_,…, *ϵ_n_* from a binding energy distribution. We associate each binding energy *ϵ*^(*i*)^ with a binding site sequence of the target TF whose energy 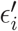 is closest to this assigned value. The sampled energy *ϵ*^(*i*)^ is then replaced with 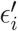, and the set of energies 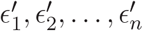 are used for all occupancy calculations in the TF-DNA binding simulation. In the simulations shown here, the energies 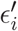 for each binding site sequence were taken from BEEML binding energy matrices (BEMs) that were fit to *in vitro* protein-binding microarray affinity measurements of different TFs [13].

Once ChIP-seq is simulated based on these binding energies, the top 10% of locations based on their read count ratios are chosen and the binding sites in each of them are used to construct the PWM of the target TF of the ChIP-seq. Since the BEMs we employed were for 10 bp long sequences, the PWMs we estimated were also 10 base pairs in length. Each term of the **W** was computed using the formula [75]

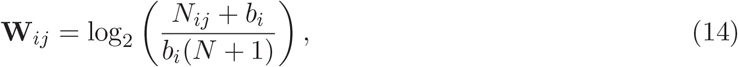

where *N_ij_* is the number of sequences with the i-th base at the j-th position, **b*_i_* is the background frequency of the i-th sequence and *N* is the number of sequences used to estimated **W**. The background frequencies are added as pseudo-counts to deal with positions where *N_ij_* is 0.

### Motif estimation in the presence of indirect binding and cooperative binding

As stated in Methods, in indirect binding, a fraction of genomic locations are bound by the ChIP target TF A while the remaining locations are bound by a second TF B, which is in turn bound to A. The binding site sequences of A are assigned to each genomic locus based on the BEM of A but we do not utilize any BEM of B to assign a binding site sequence for it. Instead, we associate with each binding energy of B a 10 bp sequence generated from a dinucleotide model of the *S. cerevisiae* genome. We chose a dinucleotide model to generate indirectly bound sequences since this has been shown to closely approximate background genomic DNA that is not bound by the target TF [61].

After ChIP-seq is simulated in the presence of indirect binding, the top 10% of genomic locations are chosen according to their read count ratio to estimate the PWM of A. The sequences used for PWM estimation are the binding sites of A from directly bound locations that fall in the top 10% and the randomly generated binding site sequences of B from the remaining indirectly bound locations.

In the case of cooperative binding, a fraction of locations is bound cooperatively by A and B while the remaining locations are independently bound by both TFs. At each genomic locus, binding sites of A and B are associated with the binding energies of each location based on their respective BEMs. When the top 10% of locations are chosen for PWM estimation, the binding sites of A from both cooperatively and independently bound locations are chosen.

### Posterior density estimation

We drew samples from the posterior binding energy distribution 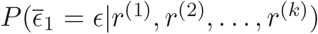 using a combination of kernel density estimation and the Metropolis-Hastings algorithm [76]. Here, 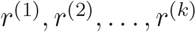 are read count ratios observed in *k* sequential replicates from a genomic locus whose true binding energy is *ϵ_T_*.

We first compute the posterior distribution when *k* = 1. From Bayes’ rule, we have

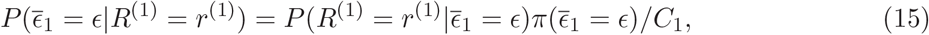

where *C*_1_ is the normalization constant 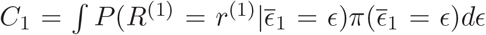 and *π* is a prior distribution, which we set to be the default binding energy distribution.

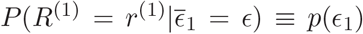 represents the probability density of read count ratios from a locus containing a single binding site with energy *ϵ*. To compute this distribution, we simulated 10^3^ replicates of ChIP-seq across *n* = 1000 locations whose binding energies *ϵ*_1_,*ϵ*_2_, …,*ϵ_n_* were sampled from the default binding energy distribution. The remaining simulation parameters of the ChIP-seq were set to their default values. This gave us 10^3^ replicates of read count ratios for each of the binding energies *ϵ*_1_, *ϵ*_2_,…,*ϵ_n_*. We used these read count ratios to compute Gaussian kernel density estimators (KDE) of *p*(*ϵ*_1_),*p*(*ϵ*_2_), …, *p*(*ϵ_n_*), which we denote as 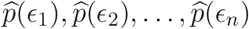. The KDE estimators were computed using the `gaussian_kde` method in the Python scipy library (v0.19.1) [77]. Given this set of KDE estimators, we can compute 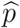 at any arbitrary *ϵ* in the range[min *ϵ*^(*i*)^, max *ϵ*^(*i*)^] using a linear interpolation between values *ϵ_j_* and *ϵ*_*j*+1_ as

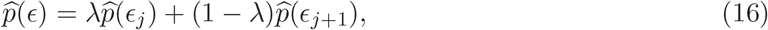

where *ϵ_j_* < *ϵ* < *ϵ*_*j*+1_ and λ = (*ϵ* – *ϵ_j_*)/(*ϵ*_*j*+1_ – *ϵ_j_*). Substituting this into equation (15), we get

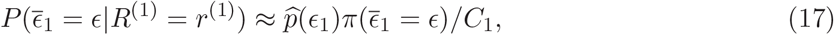

where 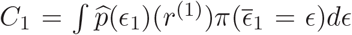. To sample from 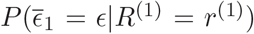, we implemented the Metropolis-Hastings algorithm [76], which is an algorithm that draws samples from a Markov chain whose stationary distribution is the posterior density from which we wish to sample.

Using the same ChIP-seq simulation parameters as used in generating the 10^3^ replicates, but with the binding energy of the first locus *ϵ*_1_ set to *ϵ_T_*, we simulated *k* replicates of ChIP-seq and store the sequence of read count ratios *r*^(1)^, *r*^(2)^,… *r*^(*k*)^ from the first locus. These are the read count ratios that are used to sequentially update the posterior binding energy estimate in equation (15).

### Metropolis-Hastings algorithm

The output of the Metropolis-Hastings algorithm is a set of samples 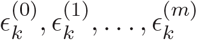 that follows a target distribution, which in this case is the posterior distribution 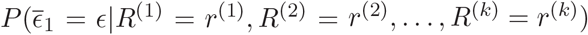. We explain the algorithm for the case when *k* = 1.

We first sample *ϵ* from a Uniform[0, 10] distribution and set 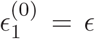, and compute 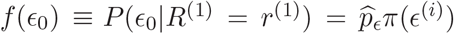. 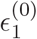 is the initial state of the Markov chain. We then sample *ϵ*′ from a proposal distribution, which we denote as 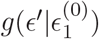, and then compute 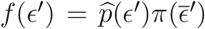 using equation (16). If 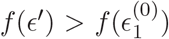, we update the current state of the sampler to 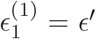. If 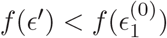, we compute a ratio

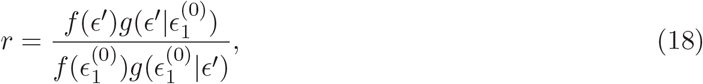

and sample a random number u from a Uniform[0,1] distribution. If *u* < *r*, we set the current state of the sampler to 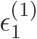 or leave it unchanged at 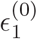 if *u* > *r*. If 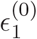 takes on values close to the values 0 or 10*k_B_T*, then the probability density 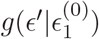 must be 0 when *ϵ*′ < 0 and *ϵ*′ > 10*k_B_T*. The choice of *g* which we use for the proposal distribution is a truncated normal distribution. The forward proposal distribution 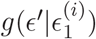, where 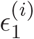 is the i-th sample drawn, is a normal distribution with mean 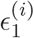 and standard deviation 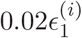 that is truncated to lie within [0, 10*k_B_T*]. The backward proposal distribution, 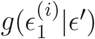 is a normal distribution with mean *ϵ*′ and standard deviation 0.2*ϵ*′ that is truncated to lie within [0, 10*k_B_T*].

We repeat this process and store the states of the sampler 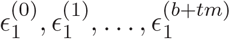, where *t* is a thinning factor that we fix at 100, *b* is the burn-in period that is set to 10000 and *m* is the number of samples to be retained, which we set at 10000. We then retain only every t-th sample beginning from the b-th sample i.e. 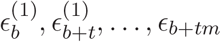, and use it to construct a Gaussian kernel density estimator of 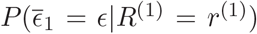, which we denote as 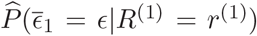. Note that in this process, the normalization constant *C*_1_ in equation (15) need not be computed.

When the read count ratio *r*^(2)^ is observed after the second replicate of ChIP-seq, we substitute 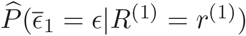 in place of the prior π in equation (15) and update the posterior 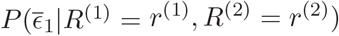 —

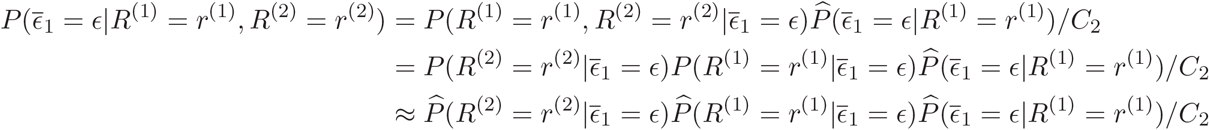

where *C*_2_ is a normalization constant. This process is repeated *k* times in order to sample from the posterior binding energy distributions 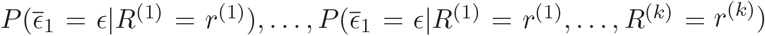

### Change in binding energy due to a single nucleotide change in a binding site

We obtained BEM models of 368 TFs from the BEEML database (http://stormo.wustl.edu/beeml/) [13]. These matrices follow the convention where the highest affinity sequence is assigned an energy of 0, with all other sequences being assigned a positive binding energy. The matrices in this database were calculated by fits of binding energy models to protein binding microarray data [13] which measured the affinity of TFs for different 10 bp long sequences. For each TF, there were two BEMs (each of dimension 4 × 10) available since the matrices were fit individually to replicates of microarray measurements, with the quality of the fit being determined by the *R*^2^ value of predictions from the matrix with measurements. For each TF, we chose the BEM that had the higher *R*^2^ value for further analysis.

For each TF, we picked the sequence whose energy was closest in value to a baseline energy that were in the range 2 – 6*k_B_T* in steps of 1*k_B_T*. For a given baseline energy, we then computed the binding energy of each of the 30 sequences that were a single mutation away from this sequence and stored the absolute values of the energy differences. We repeated this process for every TF in the database and computed the 50-th and 25-th quantiles of binding energy differences. The 50-th quantiles for the baseline energies from 2 – 6*k_B_T* were (in units of *k_B_T*) 0.24, 0.24, 0.23, 0.24, 0.24 and the 25-th quantiles were 0.76,0.72,0.7,0.7,0.71.

