## Supplement for "ChIPulate : A comprehensive ChIP-seq simulation pipeline"

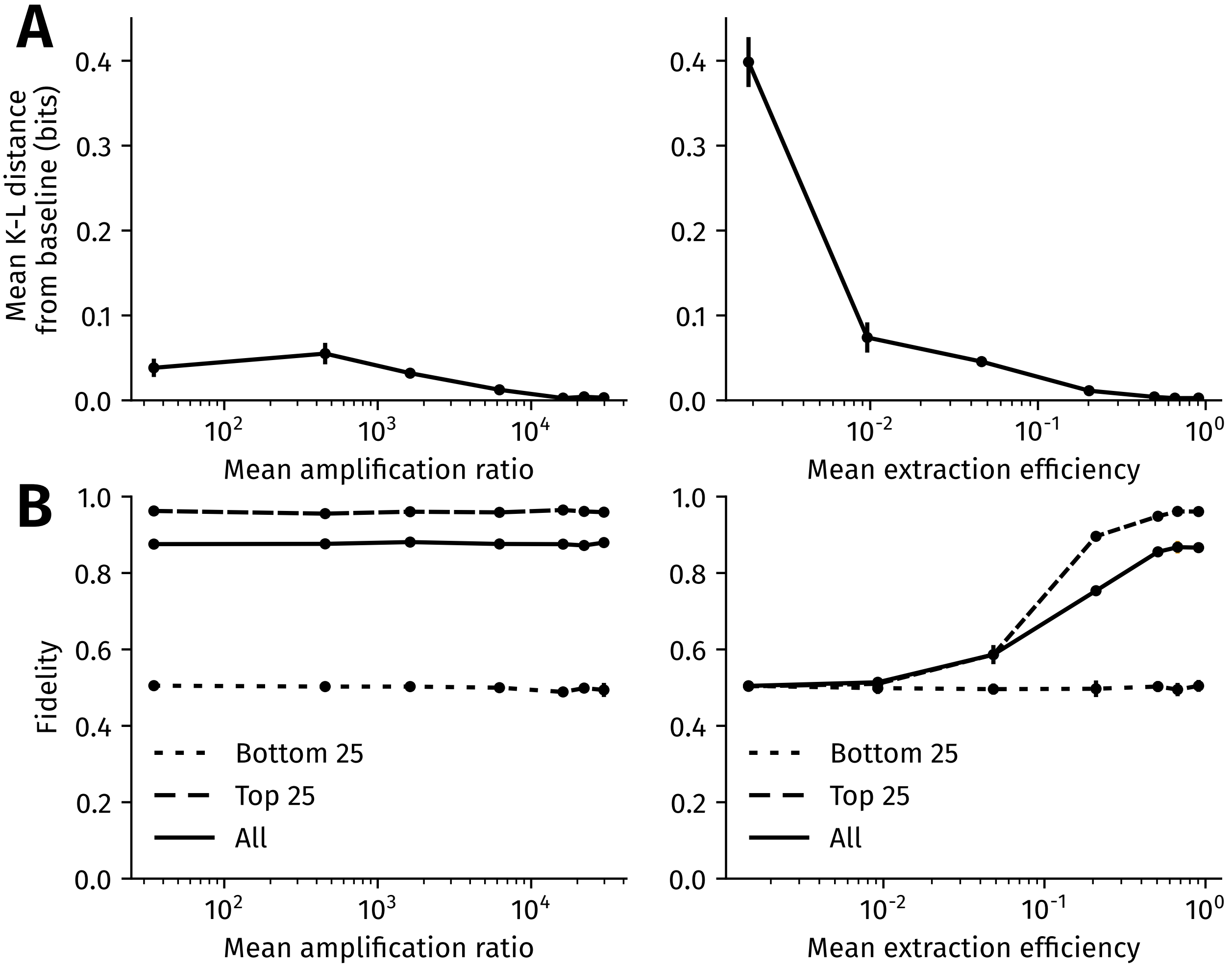


**Figure 1: Impact of power-law based genome-wide heterogeneity in extraction and PCR efficiency on motif inference and ChIP-seq fidelity.** **(A)** Impact of heterogeneity in mean amplification ratio (left panel) and extraction efficiency (right panel) on motif inference. The heterogeneity follows a power law truncated between 0 and 1. The error bars are the standard deviation in the mean K-L distance computed after PWM was estimated in 10 replicates of ChIP-seq for each mean and coefficient of variation. **(B) Impact of heterogeneity in mean amplification ratio (left panel) and extraction efficiency (right panel) on fidelity.** The y-axis is the fidelity of ChIP-seq with the mean extraction efficiency varying on the x-axis in the left panel and the mean amplification ratio varying on the x-axis in the right panel. The fidelity is calculated for all regions (orange), the top 25th percentile (blue) and bottom 25th percentile (black) of read count ratios. The error bars are the standard deviation in the estimate of fidelity, which are computed from 10 replicates of simulation for a given mean and coefficient of variation.

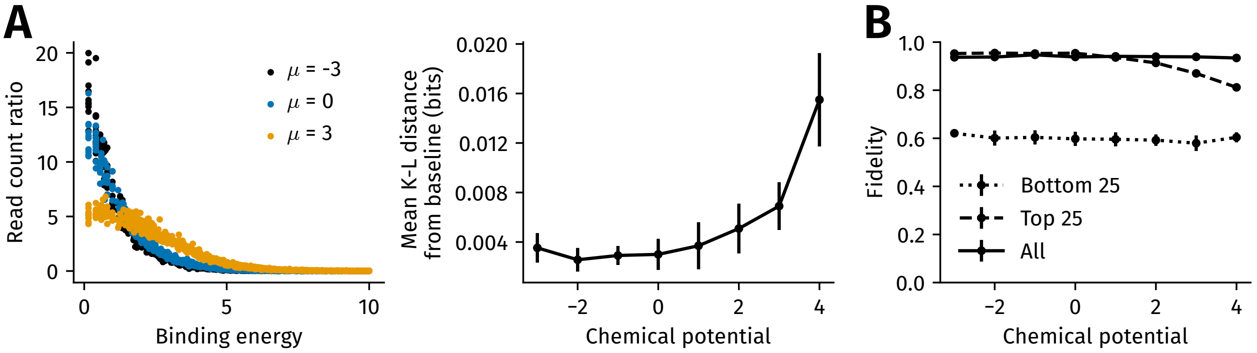


**Figure 2: Impact of chemical potential (TF concentration) on motif inference and fidelity. (A)** Left panel : Read count ratios from three different ChIP-seq simulations run with parameters set to their default values but with the chemical potential ($\mu$) set to $2k_BT$ (black), $4k_BT$ (blue) and $6k_BT$ (orange). $3k_BT$ is the default value of the chemical potential used everywhere in the main text. Right panel : The K-L distance of the derived motif of Tye7 from the baseline motif, which is derived when the chemical potential is set to 0, is shown on the y-axis. The chemical potential is varied between $1k_BT$ and $6k_BT$ on the x-axis. **(B)** The fidelity of ChIP-seq is shown on the y-axix for all regions (orange), the top 25th percentile (blue) and bottom 25th percentile (black) of read count ratios. The error bars are the standard deviation in mean K-L distance and fidelity obtained after 10 replicates of simulation.

| Domain | TF | Binding Energy Matrix | | | | | | | | | | |
|---|---|---|---|---|---|---|---|---|---|---|---|---|
| AP2 | CBF1 | A | 0.42 | 0.38 | 1.87 | 5.7 | **0.0** | 6.62 | 2.31 | 2.93 | 4.59 | 0.3 |
| | | C | 0.71 | 1.79 | 2.1 | **0.0** | 3.95 | **0.0** | 3.54 | 3.06 | 4.66 | **0.0** |
| | | G | **0.0** | **0.0** | 1.81 | 4.84 | 5.75 | 3.39 | **0.0** | 2.94 | **0.0** | 0.36 |
| | | T | 0.87 | 2.0 | **0.0** | 6.6 | 4.77 | 3.13 | 6.75 | **0.0** | 2.76 | 0.14 |
| ARID | ARID3A | A | 0.25 | 0.98 | 1.49 | 2.05 | **0.0** | **0.0** | 7.08 | 0.35 | 0.1 | 0.12 |
| | | C | 0.72 | 1.02 | 1.69 | 1.86 | 1.7 | 2.99 | 7.81 | 0.46 | **0.0** | 0.3 |
| | | G | 0.54 | 1.59 | 1.94 | 2.88 | 0.16 | 3.9 | 8.14 | 1.24 | 0.26 | **0.0** |
| | | T | **0.0** | **0.0** | **0.0** | **0.0** | 0.46 | 3.11 | **0.0** | **0.0** | 0.21 | 0.41 |
| C6 | RDS2 | A | 0.33 | 0.04 | 1.19 | 2.6 | 5.27 | 3.5 | 0.05 | **0.0** | **0.0** | 0.11 |
| | | C | **0.0** | **0.0** | 1.8 | **0.0** | 2.46 | 2.5 | 1.67 | 0.32 | 0.38 | 0.32 |
| | | G | 0.13 | 0.26 | 1.34 | 3.85 | **0.0** | **0.0** | **0.0** | 0.2 | 0.42 | 0.46 |
| | | T | **0.0** | 0.1 | **0.0** | 5.23 | 6.62 | 3.12 | 1.63 | 0.4 | 0.03 | **0.0** |
| CC | RARA | A | 0.29 | 0.38 | 3.04 | 2.06 | **0.0** | 1.57 | 2.26 | 2.08 | 0.74 | 0.23 |
| | | C | **0.0** | 0.42 | 1.4 | 2.41 | 1.83 | **0.0** | **0.0** | 0.14 | **0.0** | 0.38 |
| | | G | 0.04 | **0.0** | 4.27 | **0.0** | 2.41 | 9.49 | 8.03 | 1.27 | 0.68 | 0.19 |
| | | T | 0.26 | 0.28 | **0.0** | 3.17 | 2.44 | 5.46 | 3.45 | **0.0** | 0.13 | **0.0** |
| CH | SP4 | A | 0.73 | 0.03 | 3.14 | 1.54 | 3.87 | 7.87 | 5.63 | 0.47 | 1.68 | 0.51 |
| | | C | **0.0** | **0.0** | **0.0** | 3.37 | **0.0** | **0.0** | **0.0** | **0.0** | **0.0** | 0.46 |
| | | G | 1.21 | 3.06 | 3.97 | **0.0** | 4.7 | 4.47 | 6.34 | 3.48 | 1.68 | 1.1 |
| | | T | 0.61 | 3.1 | 3.38 | 1.44 | 3.94 | 5.67 | 1.64 | 1.56 | 0.75 | **0.0** |

| | | | | | | | | | | | | |
|---|---|---|---|---|---|---|---|---|---|---|---|---|
| ETS | ETV4 | A | **0.0** | 2.44 | 0.93 | 4.05 | 7.01 | 6.05 | 4.94 | 2.35 | 1.0 | 0.26 |
| | | C | 1.7 | **0.0** | 3.12 | 4.59 | **0.0** | **0.0** | 3.52 | 1.27 | 0.51 | **0.0** |
| | | G | 0.61 | 1.59 | 2.6 | 4.93 | 6.48 | 8.28 | **0.0** | **0.0** | 1.32 | 0.15 |
| | | T | 1.53 | 0.63 | **0.0** | **0.0** | 6.51 | 6.78 | 1.81 | 1.82 | **0.0** | 0.08 |
| GCM | GCM1 | A | **0.0** | 4.76 | 5.27 | 1.8 | 1.76 | 5.92 | **0.0** | 2.42 | 0.2 | 0.07 |
| | | C | 1.62 | **0.0** | **0.0** | **0.0** | 1.18 | **0.0** | 2.82 | 1.13 | **0.0** | 0.11 |
| | | G | 0.3 | 7.43 | 5.86 | 2.94 | **0.0** | 1.5 | 2.38 | 1.24 | 0.07 | 0.22 |
| | | T | 1.51 | 6.11 | 7.64 | 2.5 | 1.45 | 0.69 | 2.37 | **0.0** | 0.41 | **0.0** |
| HMG | LEF1 | A | 1.18 | 2.15 | 1.82 | 2.96 | 2.26 | 2.18 | **0.0** | 0.78 | 1.28 | 0.98 |
| | | C | **0.0** | **0.0** | 2.07 | 2.3 | 4.63 | 1.64 | 4.68 | 3.47 | **0.0** | 0.94 |
| | | G | 0.8 | 1.2 | 4.27 | 2.67 | 4.14 | **0.0** | 3.42 | 2.78 | 0.14 | 1.32 |
| | | T | 0.34 | 1.18 | **0.0** | **0.0** | **0.0** | 1.87 | 1.98 | **0.0** | 1.41 | **0.0** |
| LIM-homeo | LHX3 | A | **0.0** | 0.02 | 0.96 | 2.96 | **0.0** | **0.0** | 3.23 | 1.24 | **0.0** | **0.0** |
| | | C | 0.55 | 0.47 | 0.27 | 0.97 | 1.66 | 3.8 | 3.95 | 1.02 | 1.97 | 0.39 |
| | | G | 0.55 | 0.32 | 0.72 | 4.54 | 2.45 | 3.97 | 4.27 | 2.22 | 1.53 | 0.17 |
| | | T | 0.21 | **0.0** | **0.0** | **0.0** | 3.64 | 4.07 | **0.0** | **0.0** | 0.86 | 0.15 |
| POU | HDX | A | 0.05 | 0.13 | 0.12 | 0.11 | **0.0** | **0.0** | **0.0** | 1.03 | 0.82 | **0.0** |
| | | C | 0.08 | 0.12 | 0.09 | 0.79 | 0.15 | 2.16 | 6.98 | 3.09 | **0.0** | 1.61 |
| | | G | **0.0** | 0.02 | 0.16 | **0.0** | 0.28 | 0.24 | 1.07 | 1.19 | 0.71 | 0.85 |
| | | T | 0.08 | **0.0** | **0.0** | 0.82 | 0.72 | 0.67 | 1.56 | **0.0** | 1.08 | 0.77 |
| SAND | GMEB1 | A | 0.08 | 0.18 | 0.34 | 0.64 | **0.0** | 6.35 | 3.99 | 2.5 | 0.08 | **0.0** |
| | | C | 0.06 | 0.07 | 0.4 | 0.97 | 2.76 | **0.0** | 6.51 | 1.36 | **0.0** | 0.54 |
| | | G | **0.0** | 0.14 | **0.0** | 0.82 | 0.13 | 7.59 | **0.0** | 1.62 | 0.64 | 0.35 |
| | | T | 0.03 | **0.0** | 0.07 | **0.0** | 2.71 | 6.15 | 7.35 | **0.0** | 0.91 | 0.62 |
| SMAD | SMAD3 | A | **0.0** | 1.18 | 6.92 | 2.91 | 7.37 | 6.32 | 0.56 | 0.57 | **0.0** | 0.13 |
| | | C | 0.06 | 0.46 | 8.67 | 3.46 | **0.0** | 3.06 | 0.91 | 0.98 | 0.38 | 0.44 |
| | | G | 0.19 | 1.56 | **0.0** | 1.64 | 3.8 | 1.17 | **0.0** | **0.0** | 0.23 | 0.47 |
| | | T | 0.22 | **0.0** | 3.45 | **0.0** | 3.91 | **0.0** | 2.27 | 1.2 | 0.16 | **0.0** |
| T-box | EOMES | A | 0.21 | 0.53 | 2.65 | 0.55 | **0.0** | 4.48 | **0.0** | 2.04 | 1.19 | 1.26 |
| | | C | 0.39 | 0.79 | 1.39 | **0.0** | 2.26 | **0.0** | 5.48 | **0.0** | **0.0** | 0.96 |
| | | G | 0.2 | 0.64 | 2.93 | 0.62 | 0.92 | 5.87 | 1.42 | 2.18 | 1.39 | 1.97 |
| | | T | **0.0** | **0.0** | **0.0** | 1.5 | 1.17 | 3.82 | 4.06 | 3.09 | 0.93 | **0.0** |

| | | | | | | | | | | | | |
|---|---|---|---|---|---|---|---|---|---|---|---|---|
| TATA | TBP | A | **0.0** | 2.79 | **0.0** | 0.39 | **0.0** | 0.13 | 0.01 | 0.08 | 0.14 | **0.0** |
| | | C | 1.94 | 1.86 | 2.05 | 1.55 | 1.23 | 0.93 | 0.55 | **0.0** | 0.06 | 0.02 |
| | | G | 2.24 | 4.92 | 4.17 | 3.77 | 1.11 | 1.01 | 0.49 | **0.0** | 0.15 | 0.14 |
| | | T | 0.3 | **0.0** | 0.79 | **0.0** | 0.85 | **0.0** | **0.0** | 0.08 | **0.0** | 0.01 |
| bHLH | BHLHB2 | A | **0.0** | **0.0** | 1.34 | **0.0** | 4.87 | 2.92 | 2.45 | 7.16 | **0.0** | 1.08 |
| | | C | 0.24 | 0.85 | **0.0** | 2.12 | **0.0** | 3.33 | 2.73 | 5.81 | 0.43 | **0.0** |
| | | G | 0.02 | 0.23 | 3.19 | 1.54 | 3.31 | **0.0** | 2.69 | **0.0** | 2.27 | 0.47 |
| | | T | 0.14 | 0.2 | 3.23 | 1.91 | 2.43 | 5.03 | **0.0** | 8.16 | 2.17 | 0.61 |
| bHLH-ZIP | MAX | A | **0.0** | 0.35 | 1.42 | **0.0** | 4.42 | 0.61 | 2.29 | 6.58 | 0.6 | 0.15 |
| | | C | 0.39 | **0.0** | **0.0** | 2.47 | **0.0** | 2.92 | 6.73 | 9.44 | 0.36 | 0.26 |
| | | G | 0.24 | 0.27 | 2.54 | 0.88 | 2.83 | **0.0** | 4.18 | **0.0** | **0.0** | 0.23 |
| | | T | 0.42 | 0.19 | 2.74 | 1.69 | 2.44 | 5.64 | **0.0** | 3.66 | 0.41 | **0.0** |
| bHSH | TCFAP2B | A | 0.64 | 2.13 | 8.21 | 7.29 | 2.39 | 1.23 | **0.0** | 1.27 | 5.18 | 2.0 |
| | | C | **0.0** | 1.63 | **0.0** | **0.0** | 0.1 | **0.0** | 0.71 | 5.73 | 7.78 | 0.59 |
| | | G | 1.4 | **0.0** | 8.13 | 9.55 | 0.78 | 0.13 | 0.64 | **0.0** | **0.0** | **0.0** |
| | | T | 0.12 | 6.51 | 7.14 | 2.59 | **0.0** | 0.77 | 2.1 | 7.23 | 7.65 | 2.12 |
| bZIP | ATF1 | A | **0.0** | 2.87 | 2.46 | **0.0** | 2.51 | 1.34 | 0.82 | **0.0** | **0.0** | 0.19 |
| | | C | 0.87 | 4.35 | 3.48 | 6.7 | **0.0** | 3.21 | 0.75 | 0.12 | 0.62 | 0.21 |
| | | G | 0.13 | 6.18 | **0.0** | 3.33 | 2.52 | **0.0** | 1.36 | 0.3 | 0.23 | 0.17 |
| | | T | 1.2 | **0.0** | 1.07 | 1.92 | 1.1 | 1.84 | **0.0** | 0.51 | 0.37 | **0.0** |
| Fork | E2F | A | **0.0** | 0.41 | 2.88 | 3.38 | 3.52 | 7.07 | 1.03 | **0.0** | **0.0** | **0.0** |
| | | C | 0.43 | **0.0** | 0.61 | **0.0** | 6.29 | **0.0** | **0.0** | 0.28 | 0.72 | 0.42 |
| | | G | 0.12 | 0.12 | **0.0** | 8.01 | **0.0** | 1.89 | 0.72 | 0.76 | 0.87 | 0.56 |
| | | T | 0.25 | 0.8 | 3.38 | 6.51 | 1.8 | 2.75 | 1.02 | 0.38 | 0.09 | 0.15 |
| Homeo | EMX2 | A | **0.0** | 0.12 | 0.37 | 0.95 | **0.0** | **0.0** | 2.38 | 5.49 | **0.0** | 0.31 |
| | | C | 0.1 | 0.27 | **0.0** | 0.33 | 0.35 | 1.88 | 3.84 | 2.3 | 4.96 | 0.65 |
| | | G | 0.02 | **0.0** | 0.07 | 1.35 | 1.14 | 2.61 | 4.1 | 1.96 | 2.98 | **0.0** |
| | | T | 0.23 | 0.49 | 0.27 | **0.0** | 0.74 | 1.72 | **0.0** | **0.0** | 1.93 | 0.68 |
| TRP | RAP1 | A | 6.92 | 6.41 | 5.74 | **0.0** | 4.27 | **0.0** | 5.28 | **0.0** | 6.36 | 4.9 |
| | | C | **0.0** | **0.0** | **0.0** | 5.24 | 7.2 | 6.47 | **0.0** | 5.51 | **0.0** | **0.0** |
| | | G | 6.78 | 6.74 | 5.64 | 2.62 | 1.86 | 5.66 | 7.25 | 7.18 | 7.2 | 4.67 |
| | | T | 3.95 | 7.06 | 4.95 | 5.66 | **0.0** | 7.46 | 7.46 | 6.43 | 2.53 | 5.23 |

**Table 1:** The structural class of the motifs of TFs analyzed in Figure 3 and 4. Information on the structural class of the TFs was taken from the TransFac database. The binding energy matrices were taken from the BEEML database.

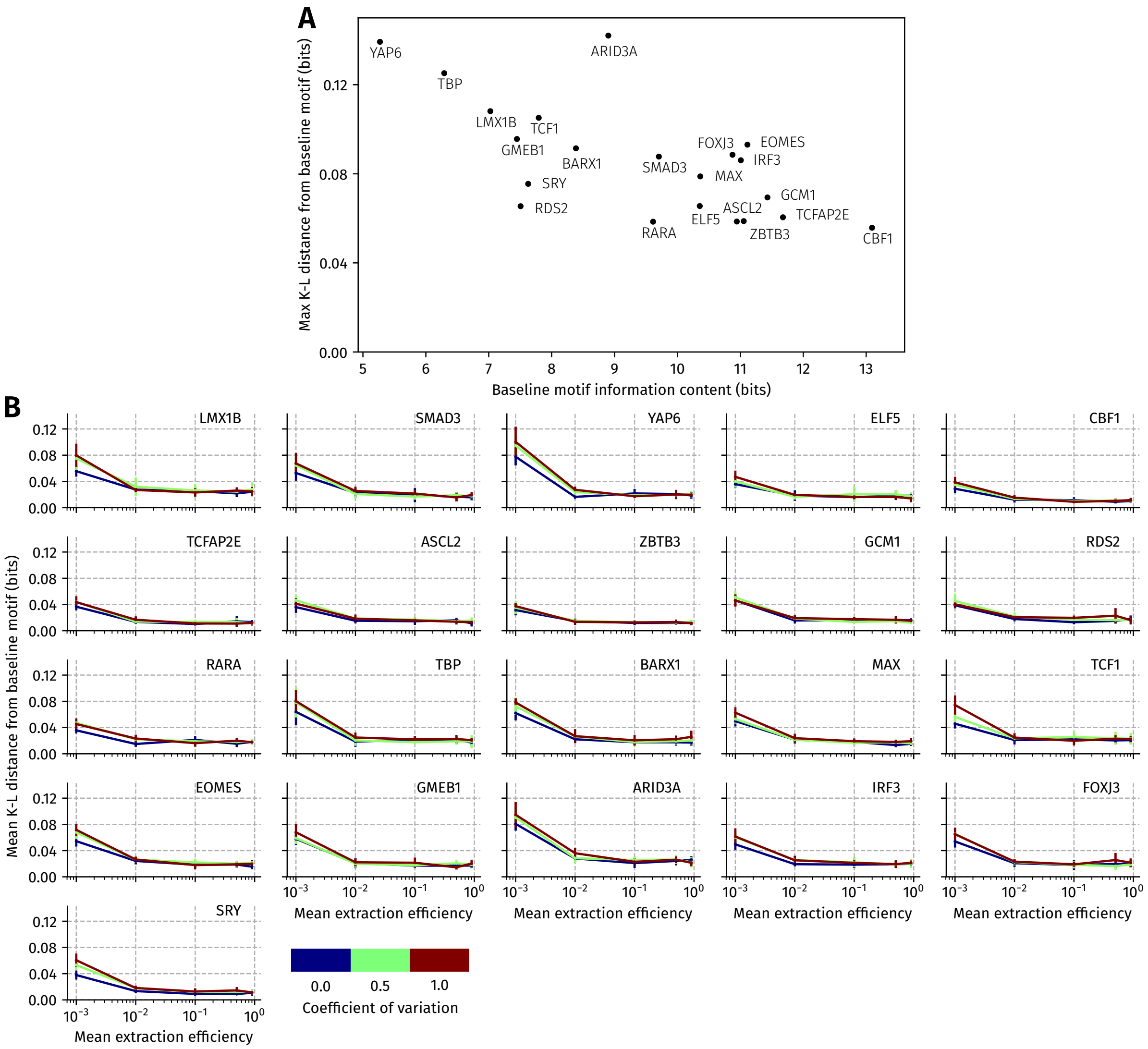


**Figure 3: The impact of normally distributed extraction heterogeneity on motif inference for different TFs.** The heterogeneity in the extraction is assumed to follow a truncated normal distribution, with the mean increasing from left to right on the x-axis in both panels. **(A) More informative TFs are less distorted by a low mean extraction efficiency.** For each TF, the maximum K-L distance between the baseline motif and the motif inferred in the presence of extraction heterogeneity is computed from the curves shown in **B**. The information content (in bits) of the baseline motif for each TF is shown on the x-axis. **(B) Dependence of K-L distance between the inferred and baseline motifs of each TF at different levels of genome-wide extraction heterogeneity.** The coefficient of variation of the truncated normal varies from 0 (no variation, in blue) to 0.5 (green) and 1.0 (brown). The error bars are the standard deviation in the mean K-L distance computed after PWM was estimated in 10 replicates of ChIP-seq for each mean and coefficient of variation. The binding energy matrices of each TF were taken from the BEEML database. The structural class of the DNA binding domains of these TFs is listed in Table 1.

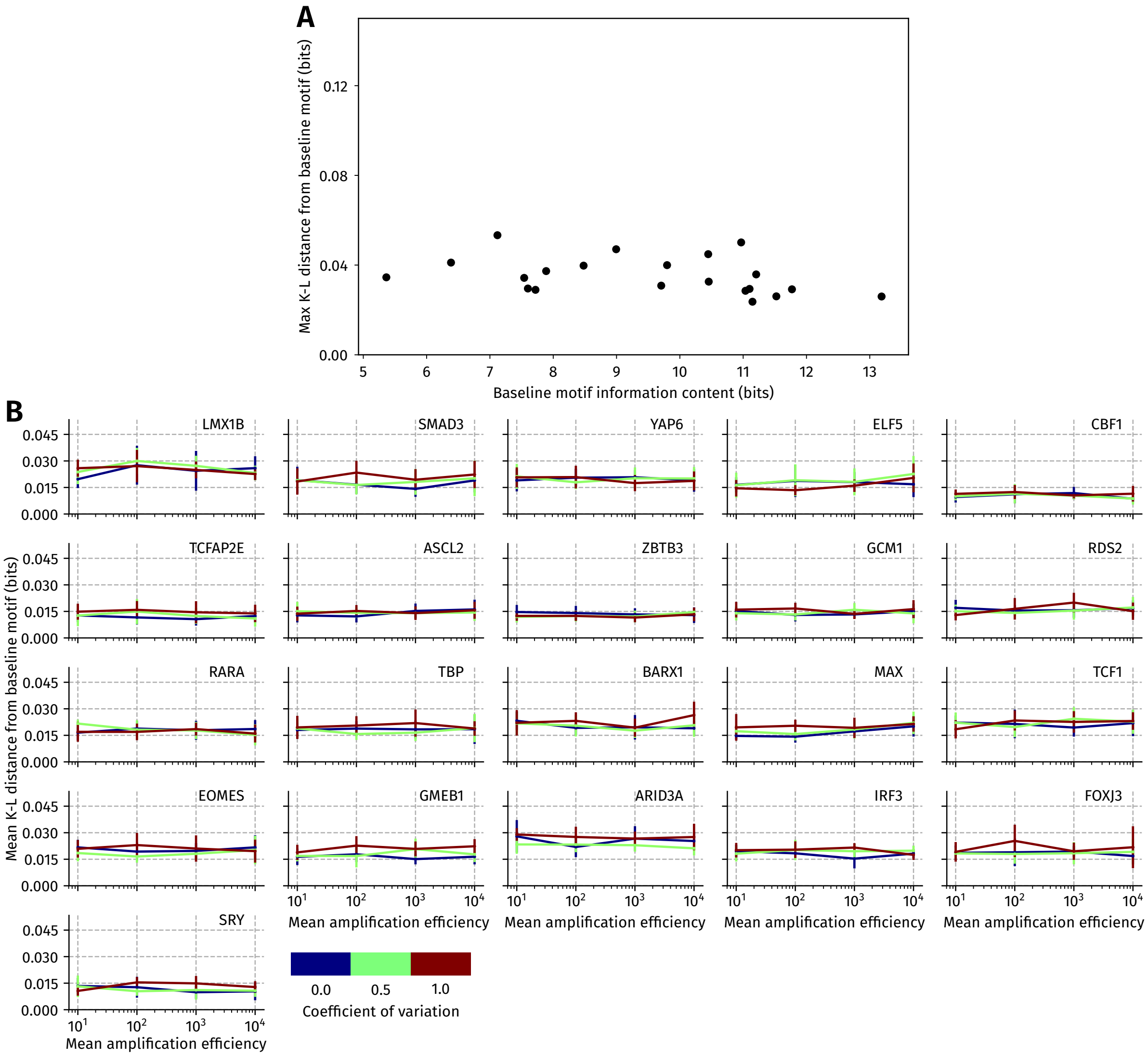


**Figure 4: The impact of normally distributed amplification ratio heterogeneity on motif inference for different TFs.** The heterogeneity in the amplification ratio is assumed to follow a truncated normal distribution, with the mean increasing from left to right on the x-axis in both panels. **(A) More informative TFs are less distorted by a low mean amplification ratio efficiency.** For each TF, the maximum K-L distance between the baseline motif and the motif inferred in the presence of amplification ratio heterogeneity is computed from the curves shown in **B**. The information content (in bits) of the baseline motif for each TF is shown on the x-axis. **(B) Dependence of K-L distance between the inferred and baseline motifs of each TF at different levels of genome-wide amplification ratio heterogeneity.** The coefficient of variation of the truncated normal varies from 0 (no variation, in blue) to 0.5 (green) and 1.0 (brown). The error bars are the standard deviation in the mean K-L distance computed after PWM was estimated in 10 replicates of ChIP-seq for each mean and coefficient of variation.

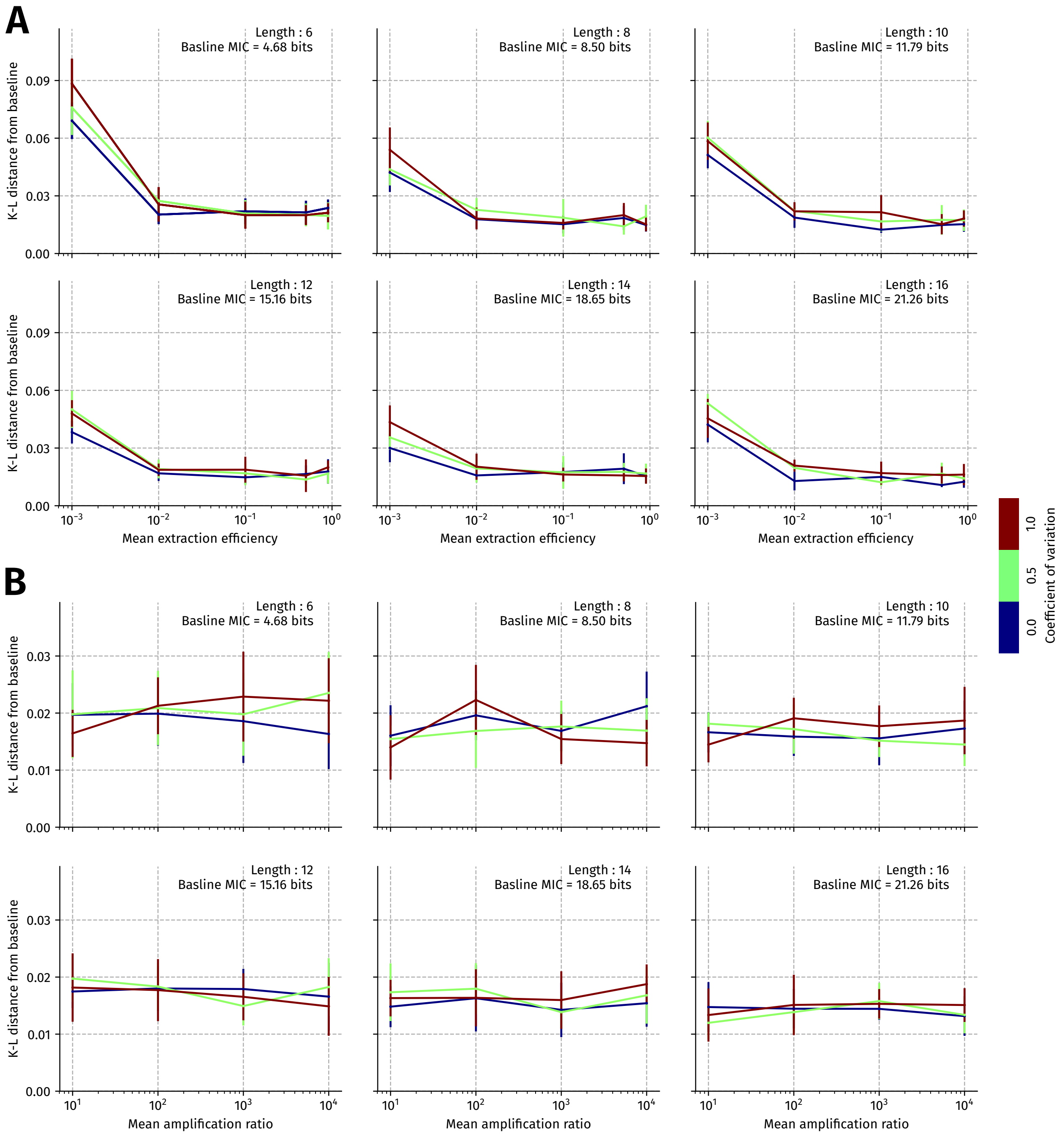


**Figure 5: Distortion of motifs of different lengths due to variation in (A) extraction efficiency, and (B) the amplification ratio.** Motifs of length 6 and 8 bp in length were generated by sub-sampling columns of the 10bp long Tye7 energy matrix, while motifs of length 12,14 and 16 were generated by first sub-sampling 2,4, and 6 columns and concatenating them to the Tye7 energy matrix. The motif information content (MIC) of the baseline motif (in bits) is shown in each panel. The extraction and amplification ratio is assumed to be normally distributed with the coefficient of variation set at either 0, 0.5 or 1. The error bars are the standard deviation in the mean K-L distance computed after the PWM was estimated in 10 replicates of ChIP-seq for each mean and coefficient of variation.

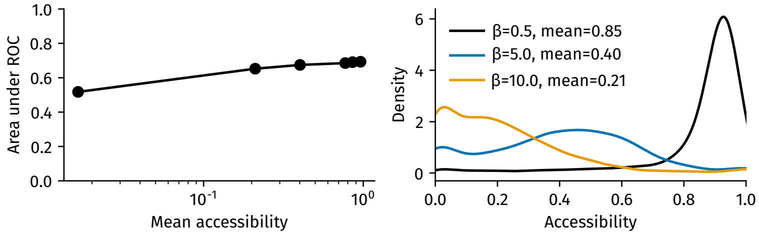


**Figure 6: The impact of chromatin accessibility on the sensitivity of ChIP-seq. Left panel :** The mean chromatin accessibility, as defined in the main text, is varied along the x-axis with the area under the ROC (auROC) on the y-axis. Equal numbers (1000) of true and false positive binding sites is set to be equal to 1000 each. The binding energy distribution of the true positive sites is set to be a power law truncated between 0 and $6k_BT$ with an exponent of 0.5, and the false positive site binding energies are distributed as a truncated power law between 0 and $6.78k_BT$ with an exponent of 0.76. This corresponds to a mean occupancy ratio of 2 between the true and false positive binding sites. Chromatin accessibility is varied by changing $\beta$ in equation 7 in the main text. **Right panel :** A Gaussian kernel density estimate of the chromatin accessibility distribution corresponding to $\beta = 0.5$ (black), $\beta = 5.0$ (blue), $\beta = 10.0$ (orange). The mean corresponding to each value is shown in the plot legend.
